# Accounting for within-species variation in continuous trait evolution on a phylogenetic network

**DOI:** 10.1101/2022.05.12.490814

**Authors:** Benjamin Teo, Jeffrey P. Rose, Paul Bastide, Cécile Ané

## Abstract

Within-species trait variation may be the result of genetic variation, environmental variation or measurement error for example. In phylogenetic comparative studies, failing to account for within-species variation has many adverse effects, such as increased error in testing hypotheses about evolutionary correlations, biased estimates of evolutionary rates, and inaccurate inference of the mode of evolution. These adverse effects were demonstrated in studies that considered a tree-like underlying phylogeny. Comparative methods on phylogenetic networks are still in their infancy. The impact of within-species variation on network-based methods has not been studied. Here, we introduce a phylogenetic linear model in which the phylogeny can be a network, to account for within-species variation in the continuous response trait assuming equal within-species variances across species. We show how inference based on the individual values can be reduced to a problem using species-level summaries, even when the within-species variance is estimated. Our method performs well under various simulation settings, and is robust when within-species variances are unequal across species. When phenotypic (within-species) correlations differ from evolutionary (between-species) correlations, estimates of evolutionary coefficients are pulled towards the phenotypic coefficients, for all methods we tested. Also, evolutionary rates are either underestimated or overestimated, depending on the mismatch between phenotypic and evolutionary relationships. We applied our method to morphological and geographical data from *Polemonium*. We find a strong negative correlation of leaflet size with elevation, despite a positive correlation within species. Our method can explore the role of gene flow in trait evolution, by comparing the fit of a network to that of a tree. We find marginal evidence for leaflet size being affected by gene flow, and support for previous observations on the challenges of using individual continuous traits to infer inheritance weights at reticulations. Our method is freely available in the `Julia` package `PhyloNetworks`.

## 1 Introduction

Phylogenetic Comparative Methods (PCM) are used to test hypotheses about the evolution of traits, using a time-scaled phylogeny to account for shared ancestry among species. For example, we consider here whether the evolution of leaflet size was correlated with biogeography, notably elevation and latitude, in the plant genus *Polemonium*. To address this question, we need to account for the correlation between species using their phylogenetic relationships. In this work, we deal with two complications: gene flow occurred in *Polemonium* [Rose et al., 2021], and leaflet size, elevation and latitude vary greatly among individual plants within a species.

Within-species trait variation is conventionally referred to as “measurement error” [e.g. Ives et al., 2007, Silvestro et al., 2015], which is a misleading term because it is too narrow. Models for trait evolution consider the mean value of a trait across a species, but this mean is usually calculated from a sample of individuals, not from the whole population. For most traits, individuals vary within a species, so the sample mean inevitably differs from the true species mean. Within-species trait variation can be due to many factors such as genetic differences, plasticity and environmental variation within a species, variation within the lifespan of an individual, or error in the act of measurement.

When the phylogeny is a tree, failure to account for within-species trait variation can lead to increased type I error [Harmon and Losos, 2005], biased and imprecise parameter estimates [Ives et al., 2007], and model selection biased towards more parameter-rich models [Silvestro et al., 2015, Cooper et al., 2016].

The impact of ignoring within-species trait variation has not been documented when the phylogeny is a network, with reticulations that can represent events such as gene flow or hybrid speciation. This is understandable since both theory and implementation of PCMs for networks are still in their infancy [Solís-Lemus et al., 2017, Bastide et al., 2018]. However, as patterns of reticulate evolution are increasingly being tested to explain phylogenetic discordance, it is crucial that the current suite of network-PCMs be expanded to account for within-species trait variation, and that the impact of this variation on inference be quantified in the context of reticulate evolution. In addition to better accounting for phylogenetic relatedness than tree-based PCMs, PCMs on networks can address new questions. For example, we quantify here the evidence that leaflet length was influenced by gene flow.

### 1.1 Existing approaches

We restrict our attention to regression PCMs, in which a response trait *y* (such as leaflet size) is modeled as a linear function of one or more predictor traits (such as elevation and latitude):

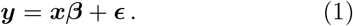

In this traditional model [Martins and Hansen, 1997], ***y*** and ***x*** contain the species means of the response and predictor traits. The residual terms in ***ϵ*** capture the phylogenetic correlation between species, based on a given phylogeny and an evolutionary model. Under the Brownian motion (BM) model on a tree, 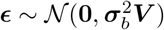 where ***V*** contains the times of shared ancestry as determined from the phylogeny, and 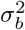 is a rate of variance accumulation [Harmon, 2019]. On a network, a hybrid’s traits are taken to be a weighted average of its parents traits (see Discussion), and this may define multiple paths from the root to a given species. In this case, ***V*** contains the expected length of the shared paths, which can be computed efficiently without having to enumerate all the paths [Bastide et al., 2018]. Beyond the BM, more flexible models can provide a continuum between low and high phylogenetic correlation like the Ornstein-Uhlenbeck model [Hansen and Martins, 1996] or Pagel’s λ [Pagel, 1999]. For these models, the phylogenetic covariance between species ***V_θ_*** is parametrized by model parameters ***θ***.

In (1), each species contributes a single value for each trait. Typically, a species’ trait value is taken to be the value from one individual, or the mean over a sample of individuals, and this sample mean is effectively treated as the true species mean. To model within-species variation, we can expand (1) to model individual values rather than species means:

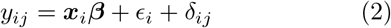

where *y_ij_* is the response of individual *j* in species *i*, ***x**_i_* contains the predictors for species *i*, and *δ_ij_* is the difference between *y_ij_* and the mean in species *i*. These *δ_ij_* values capture within-species variation in the response trait, and are typically assumed to be independent and normally distributed. Like in (1), the *ϵ_i_* values capture between-species correlation due to shared ancestry. This model was used by Ives et al. [2007], whose approach is now implemented in the R package `phytools` for instance [Revell, 2012]. In their approach, the within-species error variance is estimated separately for each species and supplied by the user (as-is or via a sample from each species). These estimated variances are then “plugged-in” as true population variances, ignoring their estimation error.

As an alternative to this “plug-in” approach, a joint estimation is used by several methods when a single observation per species is available (*j* = 1 only in (2)), such as the phylogenetic mixed model (PMM) [Lynch, 1991, Housworth et al., 2004] or `phylolm` [Ho and Ané, 2014]. With a single value per species, the covariance of the total error term *ϵ_i_* + *δ*_*i*1_ includes between-species correlations (for *ϵ*) plus an independent error variance (for *δ*) assumed equal across all species. All variance components are then estimated jointly, most often by maximum likelihood. With a single observation per species, there is no direct information about the variability within a species. Consequently, the “within-species variation” captured by this approach includes in fact any other variation that is independent across species, not already accounted for by the between-species model. On an ultrametric tree, this approach is equivalent to Pagel’s λ model [Housworth et al., 2004, Leventhal and Bonhoeffer, 2016]. With more than one observation per species and the same individuals observed across all variables (response and predictors), an alternative to regression (2) is a correlation framework to model within-species variation in all variables [Ives et al., 2007, Felsenstein, 2008]. In that approach, a model is assumed for the evolution of all variables (rather than for residuals only) and between-species relationships are represented by phylogenetic covariances between traits instead of the ***β*** coefficients. In addition, within-species relationships are represented by a multivariate phenotypic covariance matrix, whereas (2) has univariate within-species variation in the response only.

For a list of implementations that account for within-species variation, see Table 7.1 of Garamszegi [2014]. All of them assume the phylogeny to be a tree. The `Julia` package `PhyloNetworks` [Solís-Lemus et al., 2017] is currently the only available implementation for PCMs on a network. Prior to this work, `PhyloNetworks` could not account for within-species variation in the response trait other than indirectly via Pagel’s λ model.

### 1.2 Our contributions

We derived and implemented methods for model (2) along a phylogenetic network, to estimate between-species and within-species variation in the response trait, linear regression parameters, and allow for possible reticulations in the phylogeny. Our method requires that at least one species has multiple individual observations. As other regression methods based on (2), we focus on estimating the evolutionary (between-species) relationships expressed by the ***β*** coefficients. Phenotypic (within-species) relationships between traits are not modeled, as (2) uses the predictors via their species means only.

Our method differs in three main aspects from the most widely used implementations for trees. First, our method allows for one or more species to have a single observation, as is frequent in empirical data sets. This flexibility is linked to our assumption that all species share the same within-species variance of the response trait. We find that our method is robust to a violation of this assumption. Second, we do not assume that the error in sample means is perfectly known. Instead, the true within-species variance is estimated jointly with all other parameters. Finally, our implementation uses restricted maximum likelihood (REML) by default, as an alternative to maximum likelihood (ML) [Patterson and Thompson, 1971, Harville, 1974, LaMotte, 2007]. REML is known to help correct the underestimation of variance components typical of ML. For instance, Housworth et al. [2004] and Ives et al. [2007] showed that REML provides a less biased estimate of the total phenotypic variance and phylogenetic signal. Our implementation takes in individual-level data. This suggests a high computational cost. For example, with 30 species and 10 individuals per species, the input has 300 rows instead of 30 if the data were summarized by species means. As covariance matrices scale with the square of the number of rows, the cost of dealing with a much larger covariance matrix may be problematic. In this work, we show that the calculations for jointly estimating all parameters can be reduced to a computational complexity that scales with the number of species only. In fact, we show that the likelihood and restricted likelihood can be computed directly from the sample means, sample sizes, and sample standard deviations of each species. Hence, our implementation also admits this set of species-level information as input.

In the rest of the paper, we first explain the model, its assumptions, our derivations to lower the computational complexity of the (restricted) likelihood, and derivations for the reconstruction of species means. We present a thorough simulation study to assess the method’s accuracy and robustness to assumption violations, and then illustrate the method to study leaflet size evolution in *Polemonium*, whose history was shown to involve reticulation [Rose et al., 2021].

## 2 Methods

### 2.1 Model

Model (2) models within-species variation in the response variable via the *δ_ij_* term specific to individual *j* in species *i*. Let *n* be the number of species. We assume that we have data on *m_i_* individuals from species *i*, and that *m_i_* ≥ 2 for at least one species. We focus on the evolutionary correlation between the response and predictor(s), that is, the correlation over evolutionary time between the response and predictor species means. However, phenotypic (within-species) correlation can differ considerably from evolutionary correlation [Felsenstein, 1988, Goolsby et al., 2017]. For example, longevity tends to increase with body mass across species, but decrease with body mass within a species [Garamszegi, 2014]. To capture evolutionary correlations specifically, (2) uses the predictors’ species means, ignoring within-species variation for predictors.

We assume a time-scaled phylogeny including the n species of interest. If the phylogeneny is a network (also called admixture graph), each reticulation appears as a node with multiple parent edges, to represent an admixed population with genetic material from multiple parental lineages. The population inherits from each parent a proportion of genes, and inheritance proportions are assumed to be known. This punctual event is a simplified model for processes such as hybridization, horizontal gene transfer or gene flow, that can happen over a period of time [Huang et al., 2022].

For trait evolution, we assume a Gaussian model for the species-level residuals, such as Brownian motion (BM), Ornstein-Uhlenbeck (OU), or Pagel’s λ (Pλ). For now the OU model is not implemented in `PhyloNetworks`, though the subsequent derivations would equally apply. From this model and the phylogeny, we get the *n*-by-*n* unscaled covariance matrix ***V***, which may depend on some parameters, like the selection strength *α* (or phylogenetic half-life) for the OU process. For the BM on a tree, the unscaled co-variance *V_ik_* between species *i* and *k* is the length of the shared path from the root to the most recent common ancestor of *i* and *k*. To extend the BM to networks, we follow Bastide et al. [2018]. At a reticulation, the mean of the admixed population is taken to be the weighted average of the parental populations’ means. The inheritance proportions are used as weights, as reasonable for polygenic traits controlled by many additive genes. Under this model, ***V*** can be calculated in linear-time with a single traversal of the network [Bastide et al., 2018].

The evolutionary relationships are captured by a model on the true species means:

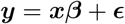

where ***y*** is the vector of the *n* true but unobserved species means for the response trait, ***x*** is an *n* × *p* matrix of predictors (including a column of ones for the intercept), ***β*** is the vector of the *p* regression coefficients, and ***ϵ*** are species-level residuals, assumed to be phylogenetically correlated: 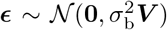. Under a BM, 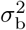 is the variance rate for the between-species residuals.

But ***y*** is unobserved. Instead, we observe a larger vector ***Y*** containing the response trait of sampled individuals. With *N* = *m*_1_ + ⋯ + *m_n_* individuals total, ***Y*** is a vector of length *N*, built by stacking the values from each species above one another. We can similarly stack the *δ_ij_* values from (2) into a vector **Δ** of length *N*, starting with the *m*_1_ values from species *i* = 1 followed by the *m*_2_ values from species *i* = 2 and so on. We can then write (2) in matrix form as follows:

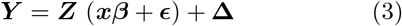

where ***Z*** is the *N* × *n* model matrix that lifts a vector of species values into a vector of individual values, by repeating the species *i* value mi times. Namely, ***Z*** is made of *n* blocks stacked above one another, with block *i* of size *m_i_* × *n*, filled with zeros except for column *i* filled with ones. More specifically, *Z_k,i_* = 1 if *m_i_* + ⋯ + *m*_*i*–1_ < *k* ≤ *m_i_* + ⋯ + *m_i_* and *Z_k,i_* = 0 otherwise.

Like earlier, we assume that the deviations from the linear relationship are phylogenetically correlated at the species-level: 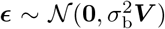. We further assume that the added within-species variation is independent with a common within-species variance 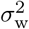: 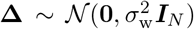 where ***I***_*N*_ is the *N* × *N* identity matrix. We can then write the full *N* × *N* covariance matrix of the total residual ***Zϵ*** + **Δ** as

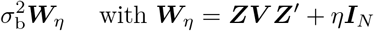

where 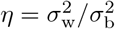.

### 2.2 Parameter estimation

If we knew *η* and any evolutionary parameters for ***V***, then (3) would be a standard linear model with known covariance and the following generalized least squares estimator for ***β***:

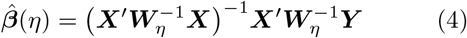

where ***X*** = ***Zx***. The above expression is rather unwieldy since it involves inverting and multiplying the large *N* × *N* matrix ***W***_*η*_. Fortunately, we show in appendix A that this expression can be simplified to:

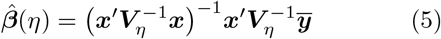

where 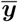 are the observed species means of the response trait and ***V***_*η*_ is *n* × *n* (much smaller than ***W***_*η*_) given by

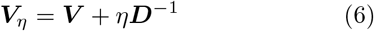

where ***D*** is the *n* × *n* diagonal matrix with the sample sizes on its diagonal: *D_ii_* = *m_i_* Note that ***V***_0_ = ***V*** corresponds to no within-species variation.

The estimation of the variance components 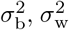 (hence their ratio *η*) and any parameters in ***V*** can be done via ML or REML. This is done by optimizing the corresponding likelihood criterion (twice the negative log likelihood) as a two-step approach. First, we fix *η* (and any parameters for ***V***) and optimize the other parameters, to obtain the *profile* criterion. In appendix A, we show that this profile criterion can be expressed using the smaller matrix ***V***_*η*_ instead of the larger matrix ***W***_*η*_ to lower the computational task:

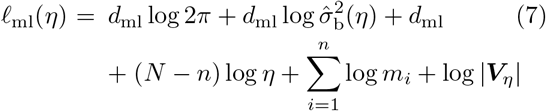

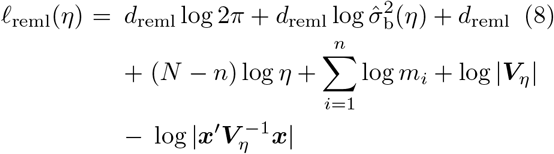

where 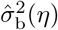, defined in (9) below, depends on the criterion via the corresponding degree of freedom: *d*_ml_ = *N* or *d*_reml_ = *N* – *p*.

We first estimate 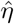 (and other parameters for ***V***) as the value that minimizes *ℓ* above. Then, we plug 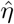 in the estimate of 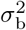 given *η*:

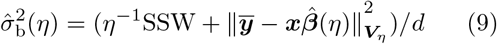

where 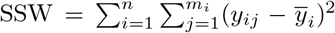 is the sum of squared residuals within species, *d* is *d*_ml_ or *d*_reml_, and 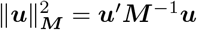. Finally, we use 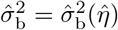 to estimate the within-species variance 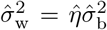, and plug 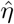 in (5) to estimate the regression coefficients.

For inference about phylogenetic coefficients *β_k_*, we implemented confidence intervals and hypothesis tests based on *n* – *p* degrees of freedom for 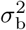. These are approximate, because *η* needs to be estimated (see appendix B). For more general model comparisons, we also implemented likelihood ratio tests.

### 2.3 Species means reconstruction

Conditional on our estimate of *η* (and of other parameters for ***V***), we can use our model to estimate the true species mean for any species in the phylogeny. For ancestral species, this task is traditionally called “ancestral state reconstruction”. This task also applies to extant species, to predict their mean based on their predictor values and data from closely related species.

Recall that ***y*** and 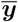 denote the true (unobserved) and the observed means, for the species with data. We further consider the true means ***y***_0_ of a set of species for which a prediction is desired. This set may include ancestral species, extant species with missing response data, or species with observed data. We assume that we know their predictor values, which we call ***x***_0_. For ancestral species, this is a very strong assumption, although it is reasonable if the predictor set is limited to the intercept column of ones or to discrete predictors that evolve sufficiently slowly for a reliable prediction in clades without variation. Another caveat, if using more than an intercept, is that the evolutionary regression (3) fitted to present-day species may not apply to past species. For example, consider a predictor *X* evolving according to a BM and a response *Y* adapting to it via an OU process with optimum *b*_0_ + *b*_1_ *X* that varies over time, as *X* evolves. Due to the lag time for adaptation, the “evolutionary slope” for *X* in ***β*** is attenuated from the “optimal slope” *b*_1_, by a factor that depends on the strength of adaptation and the height of the phylogeny [Hansen et al., 2008]. As this attenuation depends on the time from the root, inertia affects ancestral species more than present-day species under this BM-OU model, and extrapolating our regression model to ancestral states should be taken cautiously. We recommend using an intercept only for predicting ancestral states. Using other predictors should be limited to predict the mean of present-day species, or when there is evidence of fast adaptation and reliable knowledge of the predictors ancestral states.

Based on our model (3) we have that:

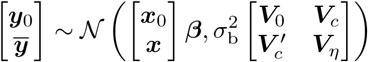

where ***V***_*η*_ is given in (6), ***V***_0_ is the phylogenetic co-variance among species for which prediction is sought, and ***V***_*c*_ is the cross-covariance of the true means, between the set of species to predict and the set of species with data. Given knowledge of 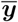, the conditional distribution of ***y***_0_ is also Gaussian with mean

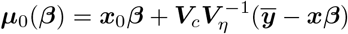

and (co)variance, or prediction error variance:

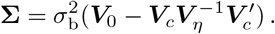

Conditional on *η* and parameters for ***V***, 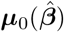 is the best linear unbiased predictor for ***y***_0_. The prediction error 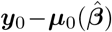 has variance equal to **∑** plus an extra term due to estimating ***β*** [Christensen, 2001] given in appendix C.1. If this prediction variance is *ψ_i_* for species *i* (which may be an ancestral or extant), then an approximate prediction interval for the true mean of that species is 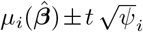 where *t* is the quantile corresponding to the desired confidence level, from the T-distribution with degree of freedom associated with 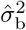 (see appendix C.2).

We note that if we have data for species *i*, then 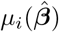 is not necessarily equal to the sample mean 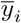. This is because the prediction is influenced by data from closely related species. Appendix C.3 illustrates this on a simple 3-species example. If many individuals are observed for a given species, then the prediction of the true mean for that species is very close to its sample mean. If few individuals are observed instead, the predicted mean is also influenced by the linear relationship with the predictors for that species, and by observations from closely related species.

### 2.4 Within-species variation in predictors

The model described above ignores within-species variation for predictors, to focus on their evolutionary relationship with the response trait. Indeed, the evolutionary (between-species) and phenotypic (within-species) relationships can be different. At the extreme, two traits can be negatively correlated within each individuals species, yet positively correlated evolutionarily [Garamszegi, 2014, Fig. 7.2]. However, if the phenotypic and evolutionary relationships are similar, then some information about this common relationship is lost when ignoring within-species (phenotypic) variation in predictors.

If one is willing to assume that the regression coefficients ***β*** are shared within and between species and if there is within-species variation in one or more predictors, then it is appropriate to consider the individual values for each predictor without summarizing the predictor data to a single average value per species. Accordingly, we consider the following model:

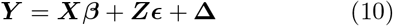

where ***Z, ϵ*** and **Δ** are as before. Here, the matrix of predictors ***X*** contains values at the individual level, where different individuals from the same species may have different values. In contrast, (3) imposes the constraint that ***X*** = ***Zx***. Note that (10) has the same parameters as (3) and the same variance 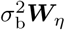 for the total residual ***Zϵ*** + **Δ**, where ***W***_*η*_ = **ZV Z**′ + *η**I**N*.

If we assume that ***V*** is from a BM and has no extra parameter, then this model is equivalent to Pagel’s λ model [Pagel, 1999] on an expanded network that has one leaf per individual, if the network is ultrametric (all leaves are equidistant from the root). Indeed, consider the expanded network constructed from the species network as follows: for each *i*, change the tip for species *i* in the original network into an internal node, then graft on this node mi external edges of length 0 (creating a polytomy of *m_i_* ≥ 3 or a degree-2 node if *m_i_* = 1) and label each new tip with an individual sampled from species *i*. This zero-length extension is similar in spirit to the approach by Felsenstein [2008]. Then, the BM covariance under this expanded network is exactly **ZVZ**′ = ***W***_0_. Bastide et al. [2018] described Pagel’s λ model on a network, which requires that the network be time-consistent (any two paths from the root to the same end node have the same length). If the distance from the root to every leaf is *h* (for height), then Pagel’s λ covariance is

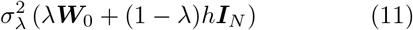

where 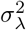 controls the total variance from the root to the tips, and λ is the proportion explained by the phylogeny. No phylogenetic signal corresponds to λ = 0 with independent observations, while λ = 1 corresponds to the BM. The variance from (10) equals that from Pagel’s λ in (11) if we reparametrize the variance components as follows: 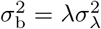 and 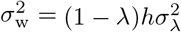, hence *η* = *h*(1 – λ)/λ.

In practice, we can fit (10) by expanding the network and using the routine developed by Bastide et al. [2018] under Pagel’s λ (which we expanded to allow for the REML criterion), then re-expressing the variance parameters in terms of between and within-species variances.

Since (10) is used with a BM model, and the coefficients ***β*** are assumed to apply both between species and within species, this model corresponds to a BM with a phenotypic relationship constrained to match the evolutionary relationship. Therefore, we abbreviate this model as BM_pheno_ later.

It is worth noting that the degrees of freedom for testing hypotheses about ***β*** is larger in model (10) than (3), because ***β*** is an individual-level parameter in (10) as opposed to a species-level parameter in (3). Intuitively, (10) makes a stronger assumption with respect to ***β*** and accordingly, it allows for a more powerful statistical test. Note that in both models, these tests are only approximate because the variance ratio *η* is estimated. Tools for mixed linear models, such as Satterthwaite’s or Kenward-Roger’s approximation [see e.g. `R` package `lmerTest`, Kuznetsova et al., 2017] or bootstrap approaches could provide more accurate confidence intervals for fixed effects and variance parameters.

### 2.5 Simulations

To quantify the performance of our method and its robustness to assumptions, we used `PhyloNetworks` to simulate trait data on a network with 3 reticulations. We used the 17-taxon network on the flowering plant genus *Polemonium* estimated by Rose et al. [2021]. We calibrated it following the approach described in Bastide et al. [2018], to obtain branch lengths proportional to time instead of branch lengths in coalescent units. The resultant network is shown in Figure 1.

**Figure 1:**
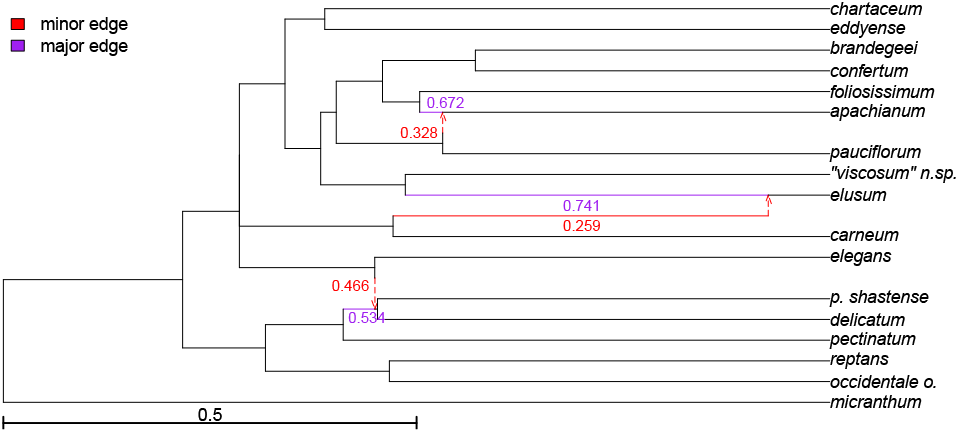
Calibrated 17-taxon SNaQ network. Edge lengths are normalized so that the network height is 1. The dotted vertical components of minor edges indicate the destination of gene flow, and do not contribute to length. Hybrid edges are labelled with their inheritance weights. The *major tree* of the network is obtained by deleting the minor (red) edges and setting the weights for the major (purple) edges to 1. The *minor tree* is obtained by deleting the major edges and setting the weights of the minor edges to 1 (e.g. *P. elusum* is sister to *P. carneum*, not *P. “viscosum” n.sp*., in the minor tree).

We describe here the most general form of our simulation model, which allows for model violation via within-species variation in the predictor and possible phenotypic correlation. Since the simulated phenotypic and evolutionary correlations may differ, our simulation model is similar to the PMM [Lynch, 1991], which has separate trait covariances for the heritable and non-heritable components (but uses a single value per species).

We simulated one predictor *X* with a BM with variance rate 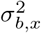 and within-species variance 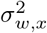:

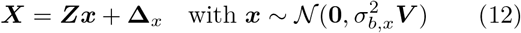

and with 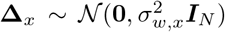. We then simulated the response *Y* as a linear function of the true species mean of *X*, an additional phylogenetic component between species, and within-species variation possibly correlated with the within-species variation in *X*:

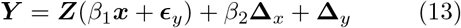

with 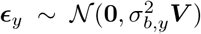 from a BM with rate 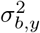 and 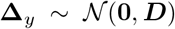 is within-species variation independent of the predictor, using 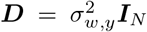 as in our estimation model, unless otherwise noted. In some simulations, we set ***D*** to be diagonal with different entries for different species, that is, unequal within-species variances. With these notations, the true species means for the response are ***y*** = *β*_1_***x*** + ***ϵ***_*y*_.

Our model (3) allows for an intercept, which we fixed to *β*_0_ = 0 in our simulations. Our model does not make any assumption on *X*, but assumes that the species means ***x*** are observed. This is the case if 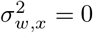, which implies that **Δ**_*x*_ = **0** and *β*_2_ becomes irrelevant. If 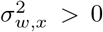, then ***x*** is unobserved, and the sample species means for *X* need to be used for estimation instead. In that case, our simulations violate the assumptions of our model. The phylogenetic and phenotypic relationships are equal if *β*_1_ = *β*_2_, as assumed in (10) by our λ-model on the expanded network.

In all of our simulations, we set 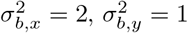 and *β*_1_ = 1. We set the sample sizes and other parameters according to various settings, as described below, and simulated 500 data sets for each combination of parameters.

We then estimated the model parameters using various methods and ML or REML. Namely, we used the BM or Pagel’s λ model that use species means, which we abbreviate as BM_n_ and Pλ_n_ (where “n” stands for “no” within-species variation). We also used model (3) under a BM, which we abbreviate as BM_y_ as it accounts for within-species variation in *Y*, but not in predictors. Finally, we used the BM_pheno_ model (10), which accounts for within-species variation in both the response and predictors but constrains the phenotypic relationship.

#### 2.5.1 Impact of ignoring within-species variation

To assess the impact of accounting for within-species variation, we used equal sample sizes *m_i_* = *m* with *m* = 3 or 8, 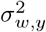 in {0.4, 0.6, 0.8}, and no model violation: 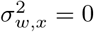. We then compared the estimates obtained with ML versus REML, and the methods that ignore or account for within-species variation.

#### 2.5.2 Impact of unequal within-species variances

Our model (3) assumes equal variances within species. To assess the robustness of our method, we used ***D*** diagonal with entry 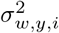 for species *i*. For each simulated data set 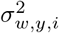, was set to a “low” value 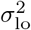 for 9 species and to a “high” value 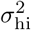 for 8 species. Species were randomly re-assigned to a low or high variance for each simulated data set. Variances 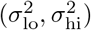 were set to (0.2, 0.2), (0.2, 0.4) or to (0.2, 0.8). We then compared the BM methods that account for within-species variation, with either ML or REML.

We again used equal sample sizes *m_i_*, = *m* with *m* = 3 or 8 and 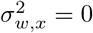.

#### 2.5.3 Impact of unequal sample sizes

In most empirical data sets, sample size variation can be substantial, with one or a few individuals from rare species, to hundreds of individuals from abundant species. To assess the impact of sample size variation, we simulated data under the same settings as in 2.5.1 except for the sample sizes *m_i_*. Like in 2.5.1, the average sample size 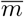 was set to either 3 or 8. For 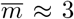, species were randomly assigned a sample size such that 5 species had *m_i_*, = 1, 6 species had *m_i_* = 3 and 6 species had *m_i_* = 5, leading to a total of 53 individuals and 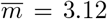. Species were re-assigned to sample sizes for each simulated data set. For 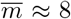, species were similarly randomly assigned such that 6 species had *m_i_* = 2, 6 species had *m_i_* = 8 and 5 species had *m_i_*, = 15, for a total of 135 individuals and 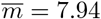.

We compared the ML and REML methods that account for within-species variation in this setting to the setting when all *m_i_* = 3 or all *m_i_* = 8 from earlier.

#### 2.5.4 Within-species variation in the predictor

Our method assumes no within-species variation in *X*. If present, this variation is ignored in practice and the sample species means are used for *X*. To assess robustness to a violation of this assumption, we simulated data as in 2.5.1 except that we set 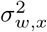 to be non-zero. Specifically, we set 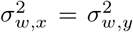. Within-species variation in *X* was uncorrelated with *Y*, that is, we set *β*_2_ = 0 to simulate the absence of phenotypic correlation. We then used the methods that ignore or account for within-species variation in the response *Y*, with ML or REML.

#### 2.5.5 Impact of phenotypic correlation

We ran the same basic settings as in 2.5.1, except that we simulated within-species variation in *X* and phenotypic correlation by setting *β*_2_ to be in { – 1,1, 2}, so that within each species *Y* is correlated with *X* with regression coefficient *β*_2_. When *β*_2_ is set to 1, the phenotypic and evolutionary coefficients are equal, as assumed by the λ-model on the expanded network. When *β*_2_ is set to –1, the phenotypic and evolutionary relationships are opposite. Also, we set *m* = 8 and 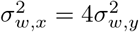 with 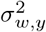 set in {0.1, 0.15, 0.2}. We used smaller values for 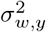 here than in 2.5.1 because the total within-species variance in *Y* is 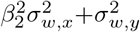, with values comparable to that in previous settings.

We then compared the estimates obtained with REML for methods ignoring within-species variation (BM_n_ and Pλ_n_), and accounting for within-species variation in *Y* (BM_y_) or in both *Y* and *X* (BM_pheno_).

### 2.6 *Polemonium* leaflet size evolution

#### 2.6.1 Objectives

We applied our method on morphological and geographical data from the flowering plant genus *Polemonium* (Polemoniaceae). *Polemonium* is widespread in North America and northern Eurasia, occurring across a broad latitudinal range from central Mexico to northern Alaska. Within its range, species of *Polemonium* can be found from sea level to the alpine zone of mountains. Vegetatively, leaves are deeply dissected (compound) into multiple leaflets. Attendant with the broad ecological amplitude of the genus is extreme variation in leaflet size across species, giving an opportunity to explore the relationship between leaflet traits and ecological predictors while accounting for phylogenetic correlation among species and trait variation within and among species.

An overall trend well-demonstrated in the ecological literature is a decreased size of vegetative structures within and across species at increasing elevations. It is thought that the wider boundary layer of large leaves (or their functional analogues) makes heat exchange more difficult and therefore large leaves are more susceptible to frost damage than small leaves [Körner et al., 1989, Wright et al., 2017]. Any relationship between morphological traits and elevation may be confounded by latitude, as high latitude communities are expected to be more ecologically similar to high elevation communities at low latitudes. Specifically, we hypothesized that leaflet size would tend to be larger in species found in low elevation, low latitude communities and smaller in species from high elevation or high latitude communities.

Because Rose et al. [2021] found evidence for reticulate evolution in *Polemonium*, we additionally investigated whether leaflet size is a trait that could have been carried along with any gene flow events. Specifically, we can test if hybridization is useful to explain residual variation, beyond the variation explained by geographical predictors.

Finally, we sought to quantify how modeling choices may impact conclusions for this dataset. As described below, choices included (1) using ML versus REML, (2) using a tree that ignores reticulation but has more taxa (therefore more data) versus using a network that better represents the phylogenetic signal but has fewer taxa, and (3) accounting for or ignoring within-taxon variation.

#### 2.6.2 *Polemonium* phylogeny

We conducted two sets of analyses, each using one of two phylogenies from Rose et al. [2021]: a 17-taxon species network inferred with SNaQ from 325 nuclear genes (Fig. 1), and a 48-taxon species tree inferred with ASTRAL from 316 genes (Fig. 2). The taxa in the network are a subset of the taxa in the tree (tips in blue in Fig. 2) because network inference methods are limited in the number of taxa they can handle. We pruned all outgroups from the ASTRAL tree. We then calibrated each phylogeny following the approach described by Bastide et al. [2018] to obtain branch lengths proportional to time. This approach uses the branch lengths in substitutions per site in the gene trees, while accounting for gene tree discordance. In total, these two trees contained up to 30 ingroup accessions that represent unique taxa (a named species or infraspecific taxon defined by morphological traits), which we will hereafter refer to as “morphs”. These morphs may or may not be monophyletic based on molecular data.

**Figure 2:**
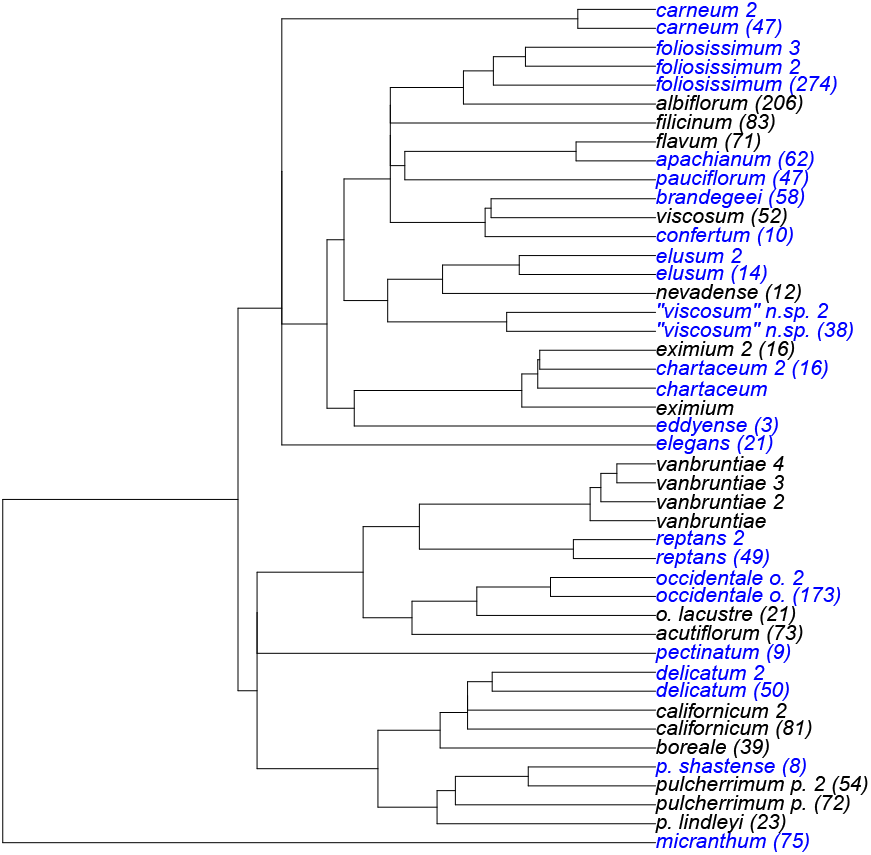
Calibrated 45-taxon *Polemonium* tree, after removing outgroups from the ASTRAL tree from Rose et al. [2021]. Edge lengths are proportional to time and normalized to a tree height of 1. The tips are labelled with their morph name, possibly with an extra index when multiple tips are from the same morph (e.g. 4 tips are from *vanbruntiae*). Morphs sampled in the network (Fig. 1) are in blue. Specimen counts for each morph are shown in parentheses, and indicate the tips retained to prune the tree to one tip per morph in Tables 1, 3 and 4.

All 17 taxa in the network corresponded to a unique morph. The tree contained 18 morphs represented by a single tip while 12 morphs were represented by two or more tips, yielding 27 tips that could not be uniquely mapped to a morph. Because our morphological data is at the species and not population level (see next) we could not assign trait values to individual tips of morphs containing multiple samples, we selected a single tip per morph in all possible ways. For 8 duplicated morphs, all tips formed a monophyletic group in the tree. Since the tree is ultra-metric, the choice of the representative tip did not affect the resulting pruned tree, so we chose one tip and deleted the others. For the 4 non-monophyletic morphs (*eximium, pulcherrimum p., chartaceum, californicum*), each one was represented by 2 tips. Because the choice of tip affects the covariance matrix, we therefore considered the 2^4^ = 16 trees obtained by choosing one of the 2 tips to represent each morph, pruning the other one from the tree. Each analysis was repeated on the 16 trees, each with a single tip per morph. One of these trees is shown in Figure 2.

To assess the signature of reticulation on leaflet evolution, we further considered two trees displayed in the 17-taxon network. First, we considered the major tree obtained by keeping all 3 major hybrid edges (which contributed a proportion of genes *γ* > 0.5 to their child hybrid node) and deleted the 3 minor hybrid edges (with γ < 0.5) from the network. Second, we considered the “minor” tree obtained by keeping the minor edges and deleting the major hybrid edges from the network (Fig. 1). The SNaQ network and the ASTRAL tree are mostly in agreement. The major tree differs from the ASTRAL tree in the placement of *P. pectinatum* and *P. pauciflorum* (Fig. 2).

#### 2.6.3 Morphology and geography data

We obtained leaflet length and width, latitude, and elevation data for all 30 *Polemonium* morphs with molecular data. Data previously published for a subset of morphs [Rose, 2021] were combined with newly generated data obtained from imaged specimens from the Consortium of Intermountain Herbaria^1^, Consortium of Pacific Northwest Herbaria^2^, Consortium of California Herbaria^3^, or loans from other herbarium collections made to JPR. Images were measured using Fiji [Schindelin et al., 2012], measuring multiple leaflets per specimen when feasible, and then averaged to obtain a specimen mean. For imaged specimens, if coordinates were present but elevation was missing, elevation was extracted from the WorldClim 2 elevation shapefile at 30-arc-second resolution [Fick and Hijmans, 2017] using the R package raster [Hijmans, 2020]. Leaflet data was obtained for between 3 to 275 specimens per morph (Fig. 2, 1757 specimens total). For the BM_y_ model, we used leaflet size from all 1757 imaged specimens, and we used the median latitude and elevation for each morph, calculated using all specimens, imaged or not (> 11000 specimens total, from 3 (latitude) and 5 (elevation) to > 1300 per morph). For the BM_pheno_ model, we only used imaged specimens for which latitude and elevation data could be extracted (997 specimens total, 2-218 per morph).

#### 2.6.4 Comparative analyses

For phylogenetic regression, we considered the following response variables: log leaflet width, log leaflet length, or log leaflet area, where the area *a* was estimated from the length *ℓ* and width *w* assuming an ellipsoid shape: *a* = *πℓw*/4. We used the natural log, a choice that impacts the interpretation of regression coefficients. We log-transformed these variables because their within-morph variance was strongly positively correlated with the mean, violating the equal-variance assumptions of our regression model. After the log transformation, the within-morph variance was stable across morphs and uncorrelated with the mean response (Fig. S1).

Using both elevation and latitude as predictors, we analyzed each measure of leaflet size using BM_y_ with REML on all phylogenies, to investigate leaflet size evolution and the signature of gene flow.

To assess the impact of model choice, we ran more extensive analyses on leaflet length, since all 3 measures showed strong positive correlation among themselves (Fig. S2). For leaflet length we used 6 methods on the full data set: ignoring (BM_n_, Pλ_n_) or accounting for (BM_y_) within-morph variation in leaflet size, using either ML or REML. We then restricted the data to specimens that had both morphological and geographical data (997 specimens), so as to use BM_pheno_. We also used BM_y_ on this data subset, to see if differences between analyses were driven by model choice or data reduction.

For each model, we recorded the coefficient estimates and their p-values, the estimated variance-component(s) and the Akaike information criterion (AIC) [Akaike, 1974]. For Pλ_n_, we conducted a likelihood ratio test of λ = 1, by comparing Pλ_n_ to the simpler BM_n_ model.

To study the impact of within-morph variation, we repeated the above analyses 100 times, each time using only a subset of 3 specimens per morph, randomly sampled without replacement from each morph.

#### 2.6.5 Leaflet size reconstruction

We demonstrate reconstruction of species means using the 17-taxon *Polemonium* network, and under the BM_y_ model with REML. To assess the effects of sample size and of model predictors on the prediction of the true mean for morphs with observed data, we predicted log leaflet length for the two morphs with smallest and the greatest number of specimens, and using or ignoring elevation and latitude as predictors. To assess the impact of node age on the uncertainty of the predicted log leaflet length, we measured the length of the prediction interval at nodes of various ages: the hybrid node ancestor to *elusum*, its minor parent, and the root (Fig. 1). For predicting ancestral states, we used a model restricted to an intercept only, as recommended above.

## 3 Results

### 3.1 Simulations

#### 3.1.1 No within-species variation in the predictor

Under settings without within-species variation in *X* (sections 2.5.1 to 2.5.3), 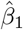 was unbiased (based on testing 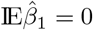 from 500 observed replicates using a t-test: *p* > 0.01 in all settings). The accuracy of 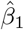 was comparable across different models, even with unequal within-species variances or varying sample sizes (Figs. 3 to 5, top).

**Figure 3:**
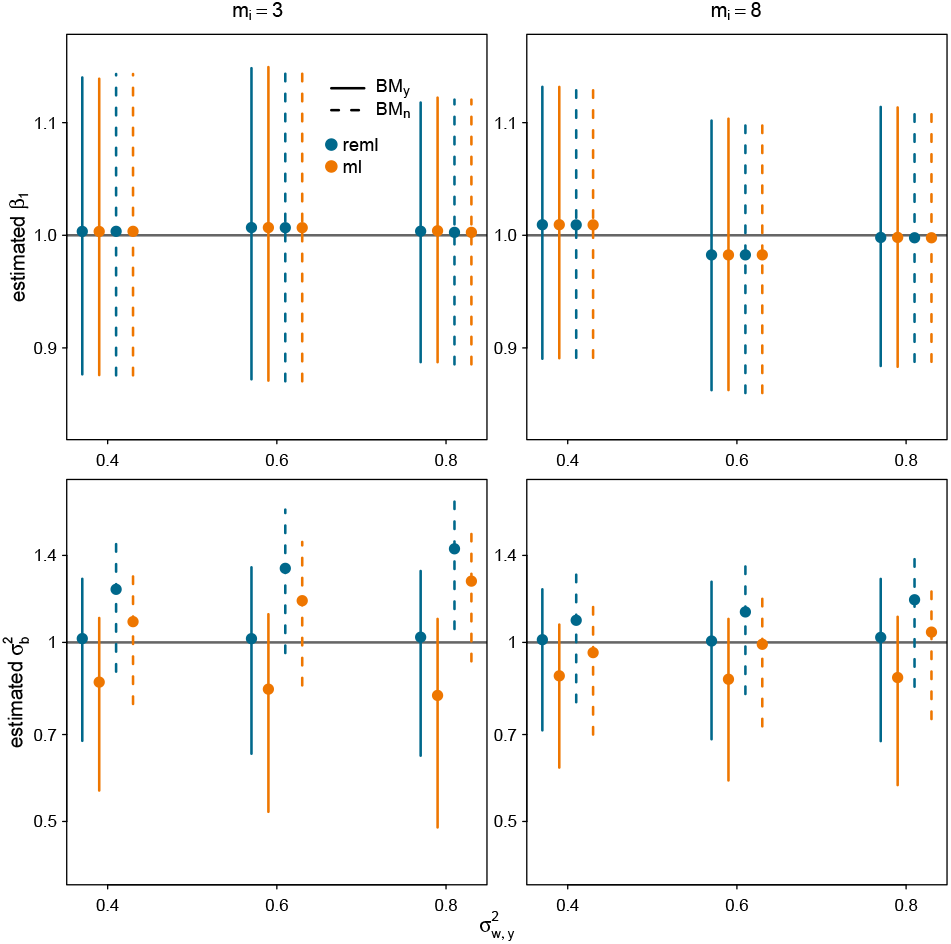
Simulations with within-species variation in *Y*, from section 2.5.1. Top: Estimated slope 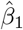. The true slope *β*_1_ = 1 is indicated by a horizontal line. Bottom: Estimated between-species variance rate on a logarithmic scale. The true value 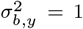 is indicated by a horizontal line. For each within-species variance 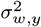 and sample size *m_i_* per species, the dots and vertical bars respectively indicate the mean and 25th-75th percentile of estimates.

**Figure 4:**
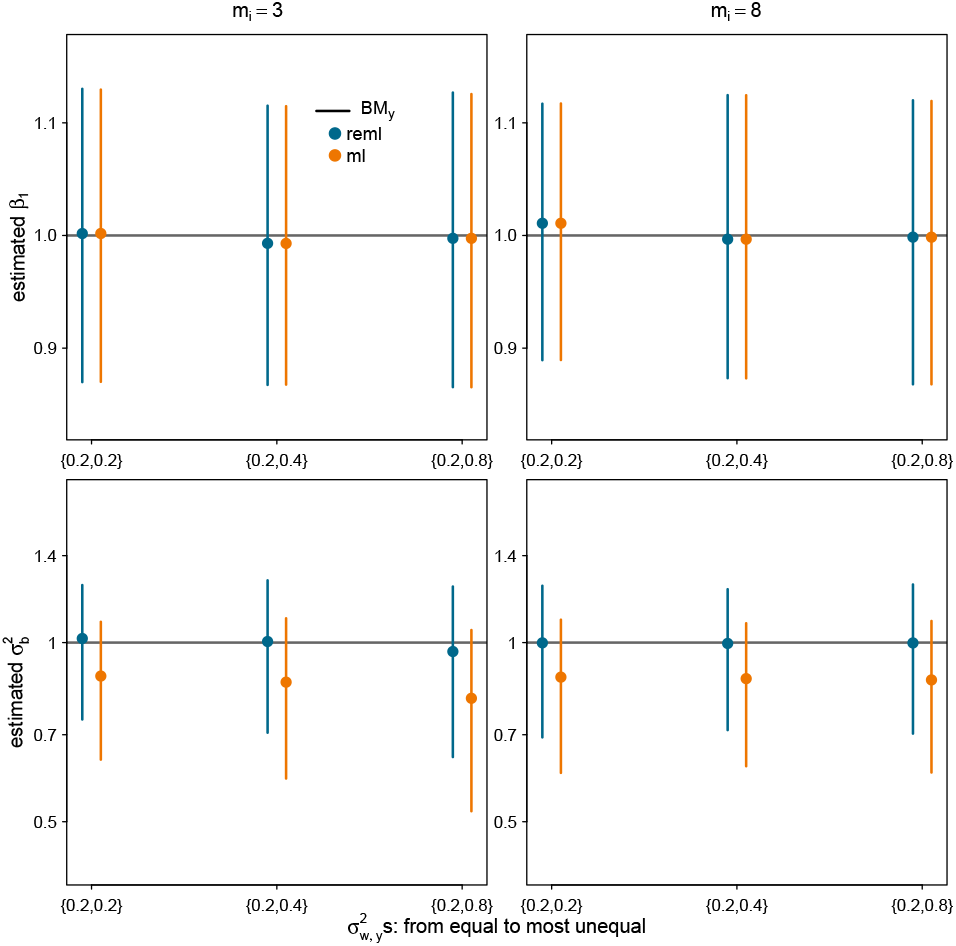
Simulations with unequal within-species variances for *Y*, from section 2.5.2. About half of species had a “low” variance and the other half had a “high” variance. Estimation used models accounting for within-species variation, but assuming equal variances.

**Figure 5:**
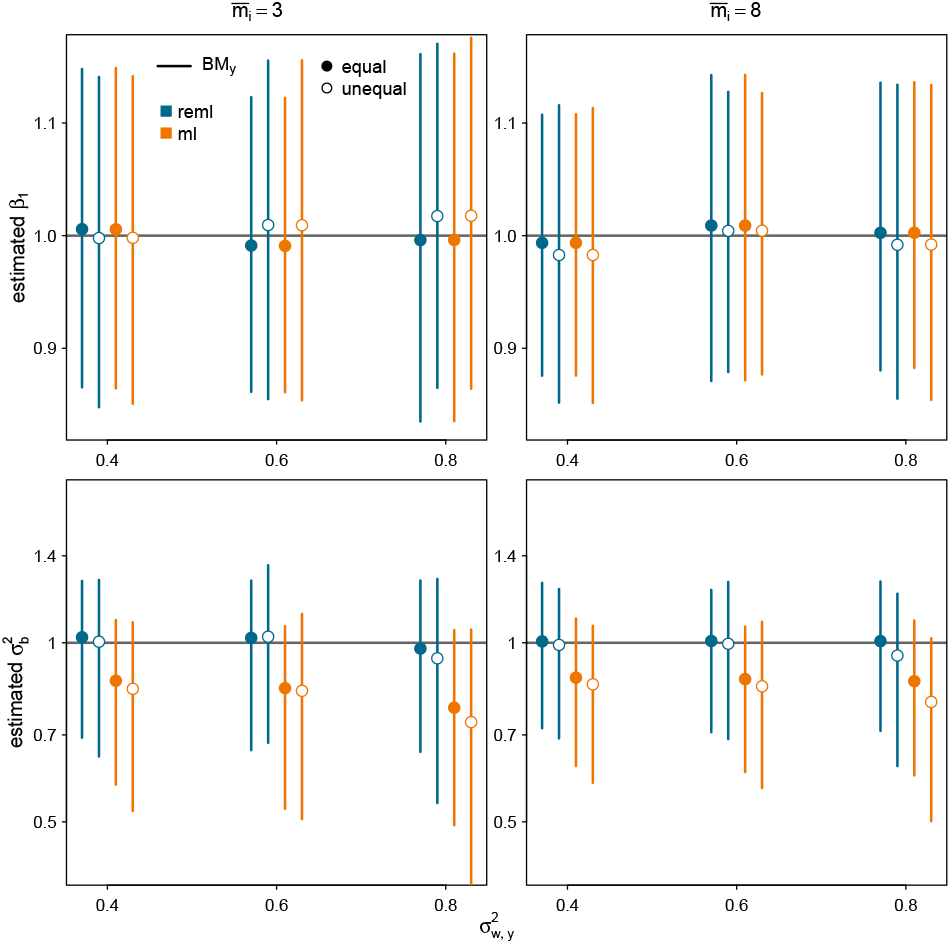
Simulations with unequal numbers of individuals per species, with average 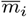 across species, from section 2.5.3. Filled dots: all species had equal sample sizes. Empty dots: species had a sample size of 1, 3 or 5 when 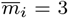, and of 2, 8, or 15 when 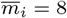.

Bias in 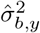 showed more sensitivity across different settings (Figs. 3 to 5, bottom). Namely, BM_y_’s estimate of 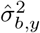 with REML was unbiased, even when sample sizes were variable or when the within-species variance varied across species. Ignoring within-species variation resulted in overestimating 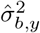, more so at smaller sample sizes. Using ML instead of REML resulted in lower estimates of 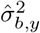, especially at smaller sample sizes. This underestimation was slightly exacerbated when sample sizes were variable.

#### 3.1.2 Within-species variation in the predictor

When within-species variation was simulated for *X* (sections 2.5.4 and 2.5.5), 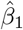 was pulled towards the true value of *β*_2_. With no phenotypic correlation, *β*_2_ = 0 so 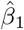 was attenuated towards 0. Since *β*_1_ = 1, all methods underestimated *β*_1_, especially so with a smaller sample size (Fig. 6, top). In general, the pull towards *β*_2_ was similar across methods that ignore within-species variation in *X* (BM_n_, Pλ_n_, and BM_y_, see Figs. 6 to 7, top) and extremely high under BM_pheno_ (Fig. 8, top).

**Figure 6:**
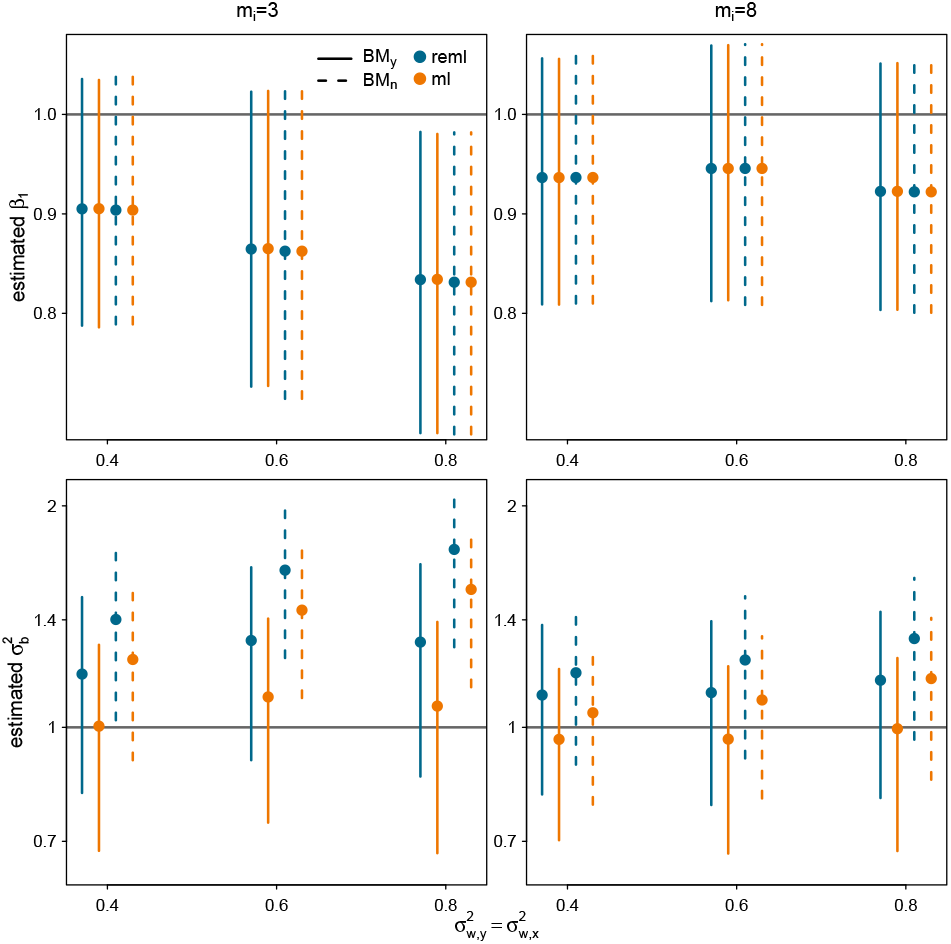
Simulations with variation within species in both *Y* and *X* but no phenotypic correlation, from section 2.5.4.

**Figure 7:**
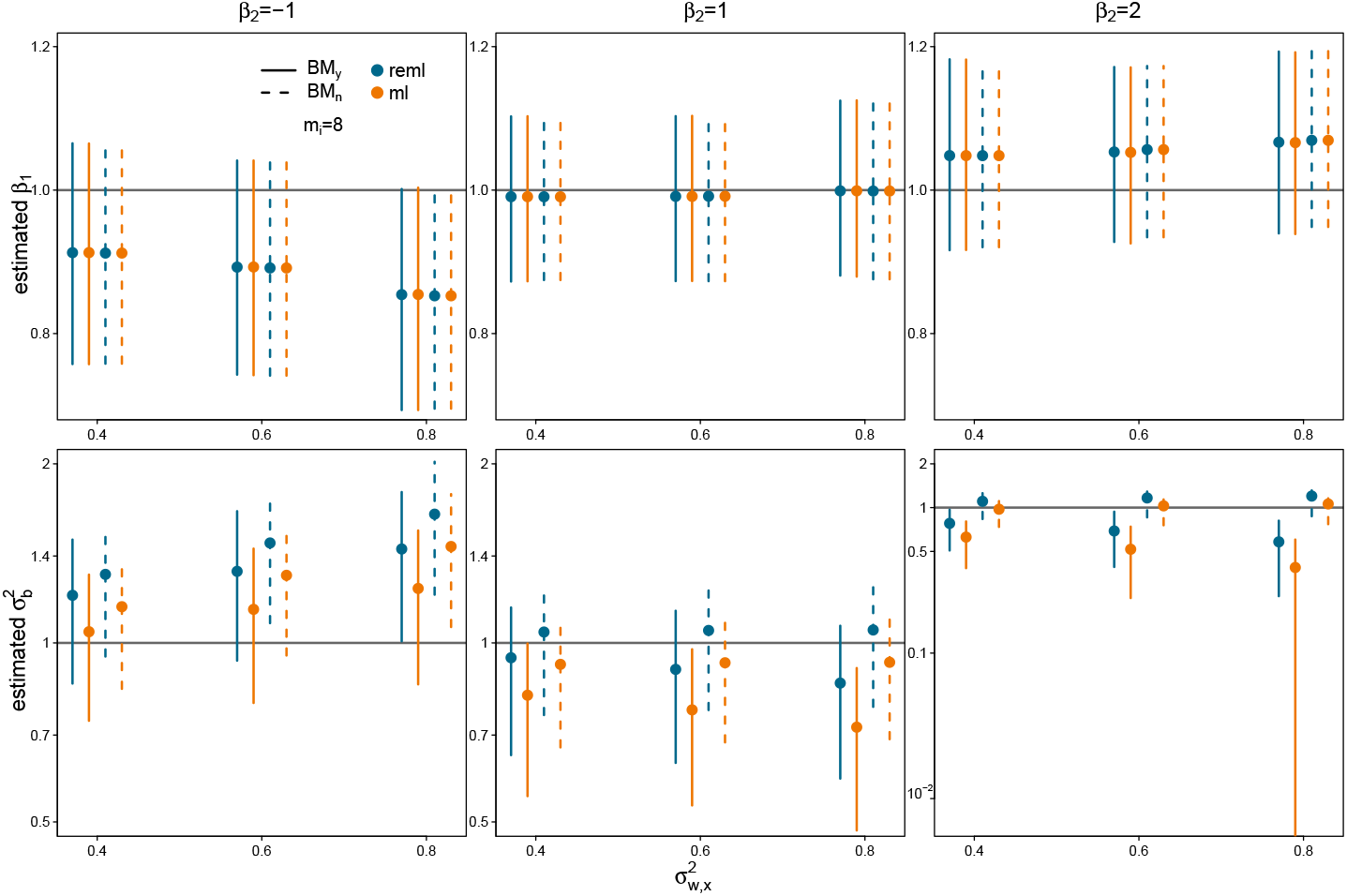
Simulations with phenotypic correlation (slope *β*_2_ within species), from section 2.5.5. Top: opposite phenotypic and evolutionary relationships (*β*_2_ = – *β*_1_). Middle: identical phenotypic and evolutionary relationships. Bottom: stronger phenotypic relationship (*β*_2_ = 2*β*_1_).

**Figure 8:**
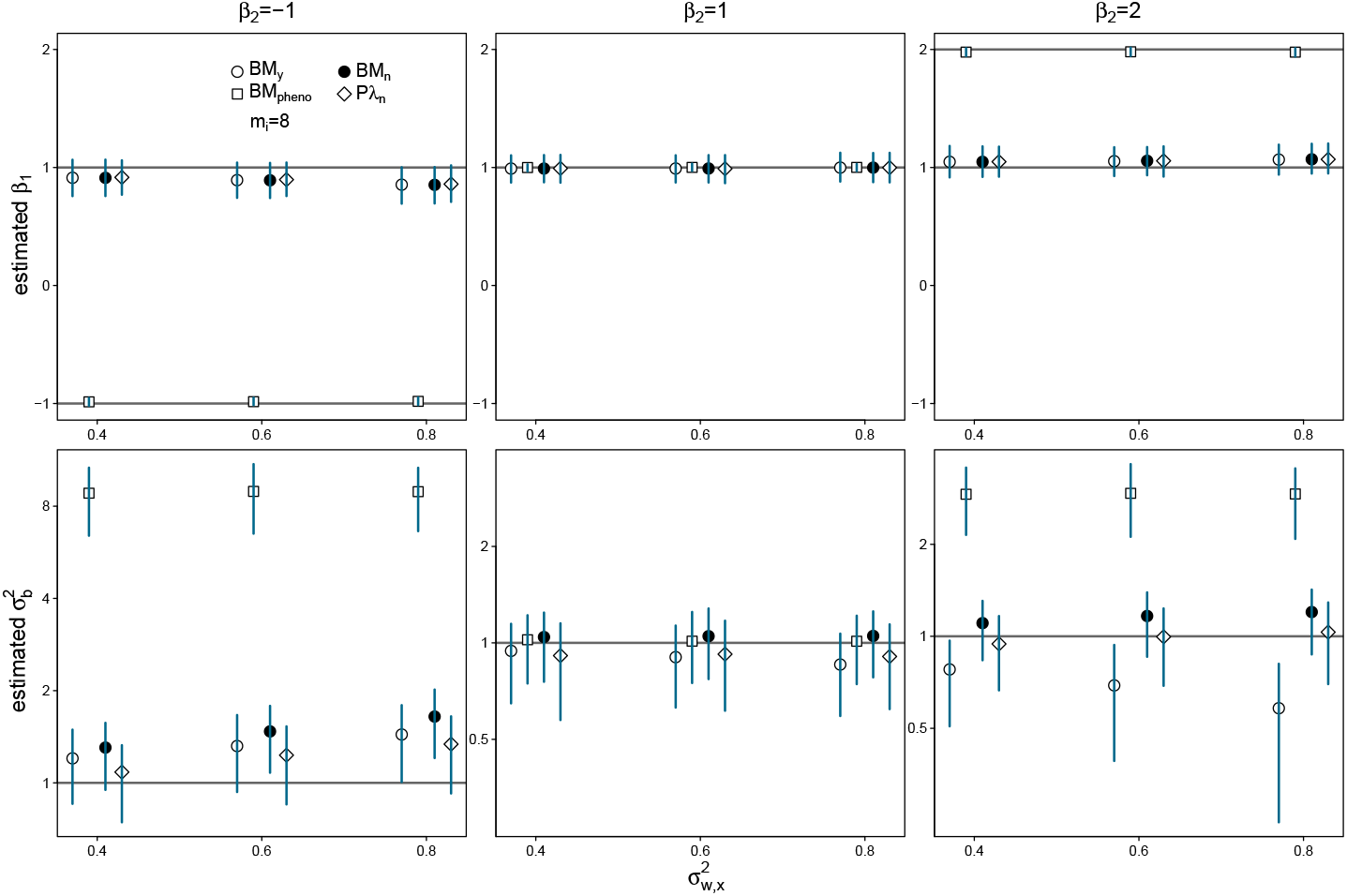
Simulations with phenotypic correlation as in Figure 7, and its impact on BM_pheno_, which assumes identical phenotypic and evolutionary relationships. The REML criterion was used for all methods: BM_n_, Pλ_n_, BM_y_ and BM_pheno_.

Like before, using ML instead of REML leads to a smaller estimate of the evolutionary variance rate 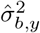, and ignoring within-species variation leads to a larger estimate (Figs. 6 to 7, bottom). However, our method BM_y_ gave a biased estimate of 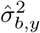 in the presence of within-species variation in *X*, with an upward bias when *β*_2_ = −1, little bias when *β*_2_ = *β*_1_ = 1 and downward bias when *β*_2_ = 2. This bias was exacerbated as 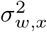 increased.

#### 3.1.3 Impact of within-species variation in *X*

To theoretically explain the bias in 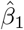and 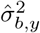 when within-species variation in *X* or phenotypic correlation is misspecified by the model, we derived the true distribution of ***Y*** conditional on the observed species means 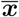, under our simulation settings. In appendix D, we show exact expressions that simplify, when *m* is large or 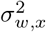 is low, to:

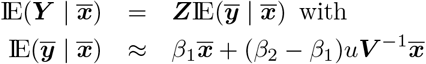

where 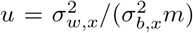. This relationship explains why our assumed evolutionary slope *β*_1_ is correctly specified if *m* → ∞ or 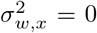 or *β*_2_ = *β*_1_. It also shows that the bias in 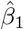 is expected to be in the direction of *β*_2_ – *β*_1_, hence the pull towards *β*_2_.

For the residual variance, appendix D shows that

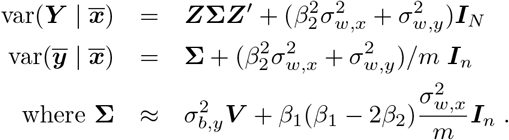

In comparison, our model BM_y_ assumes 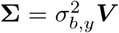. Therefore, if we focus on the diagonal terms in ***V*** (which are the largest) and their average 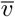, then our model expects these terms to be around 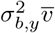, while the data will provide values around 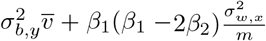. Based on these diagonal terms only, we can expect 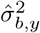 to be around 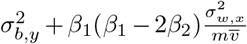, which explains a positive or negative bias, depending on how *β*_1_ compares to 2*β*_2_. If *β*_2_ = 0 for example, then we expect a positive bias (overestimation) as observed in 2.5.4 (Fig. 6, bottom). If *β*_2_ = –*β*_1_, then we also expect a positive bias. But if *β*_2_ = 2*β*_1_, then we expect a negative bias (underestimation). This is indeed what we observed in 2.5.5 (Fig. 7, bottom).

### 3.2 *Polemonium* leaflet size

#### 3.2.1 Small leaflets correlate with high elevation

The 16 ASTRAL subtrees provide extremely similar results, with parameter estimates that do not exceed 1% difference among one another (Table S1). Table 1 shows the results from one of these subtrees, selected because it gave the lowest AIC for all of the leaflet size variables.

**Table 1:**
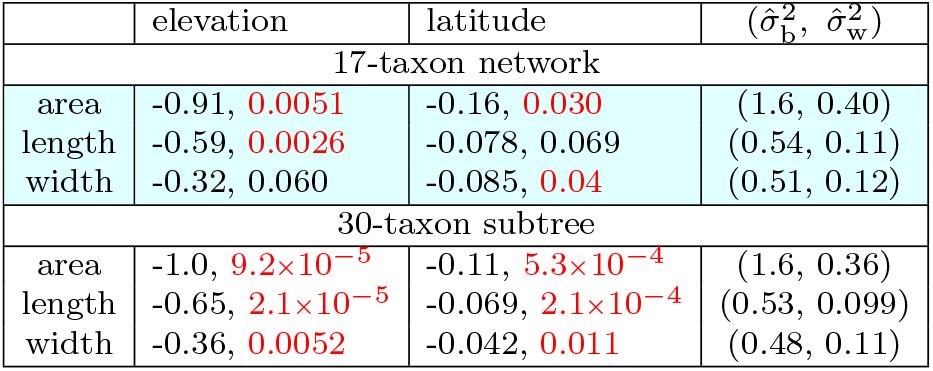
Results from fitting our model BM_y_ with REML on *Polemonium* data, to explain variation in leaflet area, length and width (each log-transformed) using elevation and latitude as predictors simultaneously. In the elevation and latitude columns, the first value is the estimated regression coefficient followed by the p-value to test that the coefficient is 0, colored in red when < 0.05.

| | elevation | latitude | $(\hat{\sigma}_b^2, \hat{\sigma}_w^2)$ |
| --- | --- | --- | --- |
| 17-taxon network |  |  |  |
| area | -0.91, 0.0051 | -0.16, 0.030 | (1.6, 0.40) |
| length | -0.59, 0.0026 | -0.078, 0.069 | (0.54, 0.11) |
| width | -0.32, 0.060 | -0.085, 0.04 | (0.51, 0.12) |
| 30-taxon subtree |  |  |  |
| area | -1.0, $9.2 \times 10^{-5}$ | -0.11, $5.3 \times 10^{-4}$ | (1.6, 0.36) |
| length | -0.65, $2.1 \times 10^{-5}$ | -0.069, $2.1 \times 10^{-4}$ | (0.53, 0.099) |
| width | -0.36, 0.0052 | -0.042, 0.011 | (0.48, 0.11) |

Elevation and latitude correlate negatively with leaflet size regardless of the measure used for leaflet size (length, width or area) or of the phylogeny (Table 1). Using the network and its 17 taxa, the elevation coefficient is negative with strong evidence for area and length, and moderate evidence only for width. The latitude coefficient is negative with moderate evidence for all 3 measures of leaflet size. Using the tree and its larger set of taxa, both elevation and latitude are negative with very strong evidence for area and length, and strong evidence for width. The p-values are smaller on the tree than on the network. This is unsurprising since the tree has almost twice the number of taxa (from 17 to 30) and specimens (from 954 to 1757). Therefore, if a true relationship exists, then the tree is expected to have more power to detect it, unless model misspecification due to using a tree causes a decrease in power (if a true relationship exists) or an increase in type-1 error (if no relationship exists). Here, the effect of phylogenetic placement is expected to be minor because the network and tree are in good agreement: the tree pruned to 17 taxa is displayed in the network, except for a small change in the position of *pectinatum*.

The coefficients are very stable across the two phylogenies. They remain negative across all three responses. The results for area are consistent with the results for length and width. On the network for instance, the elevation coefficients for log(area), log(length) and log(width) in Table 1 translate to an expected decrease in area, length and width by approximately 40%, 55% and 73% for a 1000-meter increase in elevation.^4^ These changes are consistent with one another since 0.40 ≈ 0.55 · 0.73.

The estimated variance components (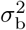 and 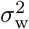) are also very stable across the two phylogenies (within 10% of each other). The ratio of within-to-between species variances, 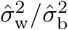 is relatively stable across measures of leaflet size (ranging from 0.18 to 0.26).

#### 3.2.2 Gene flow may explain residual variation

To test the importance of gene flow in leaflet size evolution, we compared the fit of the network to the fit of trees on the same 17-taxon data. We considered two tree models, using the “major” and the “minor” trees displayed in the network, representing the largest and the smallest proportions of the genome respectively, based on inheritance along hybrid edges in the network.

Regression coefficients estimated from these trees are fairly similar to those from the network, and the qualitative conclusions about evolutionary correlations with elevation and latitude are mostly unchanged (Table 2 and upper rows of Table 1 highlighted in cyan). The percent change in the trees’ estimates compared to the network’s estimates ranges from 0.09-13% for elevation, 0.93-11% for latitude, and 0.88-2.0% for 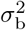.

**Table 2:** Same as Table 1, but without reticulation: using either the major or minor tree displayed in the 17-taxon network. The elevation and latitude columns contain the estimated regression coefficient, with a star (*) to indicate an associated p-value < 0.05. The last column contains the change in AIC resulting from removing reticulations: AAIC = AlC(tree) — AlC(network). The positive values indicate that the network provides a better fit than the tree in all cases.

| tree | elevation | latitude | $(\sigma_b^2, \sigma_w^2)$ | $\Delta_{AIC}$ |
| --- | --- | --- | --- | --- |
| area |  |  |  |  |
| major | -0.91* | -0.16* | (1.6, 0.40) | 0.56 |
| minor | -0.93* | -0.18* | (1.6, 0.40) | 0.28 |
| length |  |  |  |  |
| major | -0.60* | -0.078 | (0.53, 0.11) | 0.40 |
| minor | -0.56* | -0.087 | (0.54, 0.11) | 0.18 |
| width |  |  |  |  |
| major | -0.31 | -0.082 | (0.51, 0.12) | 0.87 |
| minor | -0.36* | -0.089* | (0.50, 0.12) | 0.13 |

The change in AIC from the network to a tree Δ = AIC(tree) – AIC(network) is positive regardless of which tree or response variable is used, supporting our hypothesis that gene flow explains residual variation: a reticulate network is a better representation of leaflet size evolution than a tree. However, Δ < 1 in all cases, meaning that the network is not especially helpful for explaining residual variation in the model beyond what can already be explained using either tree. Similarly, the minor tree’s AIC is better than but close to the major tree’s AIC, suggesting that the leaflet data is only marginally better explained by the minor tree than by the major tree.

#### 3.2.3 Impact of modeling choices

For log leaflet length, we explored the impact of three modeling choices: using ML instead of REML, ignoring within-species variation, and fitting BM_pheno_. We begin by addressing the first two, which involve comparing BM_y_, BM_n_ and Pλ_n_, fitted with ML and REML. The estimated coefficients for elevation and latitude were very stable across these methods, as were the magnitude of their associated p-values (Table 3). The estimated within-species variance 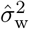 was also fairly stable across methods that estimated it. This may be due to around a third of the morphs having large sample sizes (>50 specimens). The method choice had most impact on the estimated evolutionary variance rate, 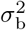.

**Table 3:** Analysis of leaflet length (log-transformed) on the full data set. BM_n_ and Pλ_n_ ignore within-species variation. In the elevation and latitude columns, the first value is the estimated coefficient. The second italicized value is the associated p-value, in red when < 0.05. The maximal possible λ (1.08 for the network and 1.14 for the subtree) may exceed 1 depending on the terminal branch lengths in the phylogeny. λ > 1 means greater phylogenetic correlation than under a BM. In the 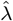 column, the first value is the estimate and the second italicized value is the p-value from the likelihood ratio test of λ = 1, which corresponds to the BM model.

| method | elevation | latitude | $\hat{\sigma}_b^2$ | $\hat{\sigma}_w^2$ | $\hat{\lambda}$ |
| --- | --- | --- | --- | --- | --- |
| 17-taxon network |  |  |  |  |  |
| BM<sub>y</sub>, REML | -0.59<br>*0.0026* | -0.078<br>*0.069* | 0.54 | 0.11 |  |
| BM<sub>y</sub>, ML | -0.59<br>*0.0012* | -0.078<br>*0.047* | 0.45 | 0.11 |  |
| BM<sub>n</sub>, REML | -0.59<br>*0.0025* | -0.078<br>*0.069* | 0.55 |  |  |
| BM<sub>n</sub>, ML | -0.59<br>*0.0025* | -0.078<br>*0.069* | 0.45 |  |  |
| Pλ<sub>n</sub>, REML | -0.59<br>*0.0024* | -0.079<br>*0.066* | 0.53 |  | 0.95<br>*0.94* |
| Pλ<sub>n</sub>, ML | -0.61<br>*0.0016* | -0.088<br>*0.047* | 0.32 |  | 0.0<br>*0.30* |
| 30-taxon subtree |  |  |  |  |  |
| BM<sub>y</sub>, REML | -0.65<br>*2.1×10<sup>-5</sup>* | -0.069<br>*2.1×10<sup>-4</sup>* | 0.53 | 0.099 |  |
| BM<sub>y</sub>, ML | -0.65<br>*9.8×10<sup>-6</sup>* | -0.069<br>*1.1×10<sup>-4</sup>* | 0.48 | 0.099 |  |
| BM<sub>n</sub>, REML | -0.65<br>*2.0×10<sup>-5</sup>* | -0.069<br>*2.1×10<sup>-4</sup>* | 0.54 |  |  |
| BM<sub>n</sub>, ML | -0.65<br>*2.0×10<sup>-5</sup>* | -0.069<br>*2.1×10<sup>-4</sup>* | 0.49 |  |  |
| Pλ<sub>n</sub>, REML | -0.64<br>*1.9×10<sup>-5</sup>* | -0.070<br>*1.9×10<sup>-4</sup>* | 0.64 |  | 1.11<br>*0.19* |
| Pλ<sub>n</sub>, ML | -0.64<br>*1.9×10<sup>-5</sup>* | -0.070<br>*1.9×10<sup>-4</sup>* | 0.57 |  | 1.11<br>*0.21* |

Using ML instead of REML caused a large decrease in 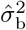 under BM_y_, by 18% on the network and 10% on the tree (Table 3). The smaller 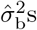 resulted in smaller standard errors for the coefficients under BM_y_, and hence smaller p-values. In our study, this moderate decrease (<0.03) in p-values changed the qualitative conclusions for latitude only. But in other studies, a decrease in p-values may result in more (or more drastic) qualitative changes in conclusions, and possibly inflated type I error rates from using ML compared to REML.

Using Pλ_n_, the choice of ML versus REML can lead to an extreme difference in the estimated λ and apparently contradictory conclusions: from no phylogenetic correlation when 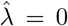 under ML, to almost full phylogenetic correlation as expected from the BM when 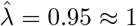 under REML. This large change occurs on the network only (Table 3), which has only about half of tree’s taxa, and is therefore less informative about phylogenetic correlation. The contradiction disappears when we use a likelihood ratio test. Using the network and either ML or REML, the likelihood is rather flat, such that there was no evidence to reject the hypothesis of a BM (λ = 1), and also no evidence to reject the lack of phylogenetic signal (λ = 0). From the larger taxon set, using either ML or REML, there was no evidence to reject λ = 1 but strong evidence to reject λ = 0. This finding supports our use of the BM model when accounting for within-species variation (BM_y_).

That 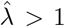 on the tree is surprising, because it means that within-species variation in leaflet length is estimated at 0. This may reflect error in the estimated tree topology or branch lengths. This may also be due to large sample sizes, leading to small standard errors in the estimated species means: 13 of the 30 morphs in the tree have over 50 specimens. On the network, fewer morphs (5 out of 17) have > 50 specimens, and 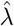 is smaller. In data sets with small sample sizes, within-species variation causes greater error in species means, and we expect Pagel’s λ to capture within-species variation as part of the non-phylogenetic signal. Indeed, 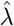 was smaller when sub-setting our data with only 3 specimens per morph. For instance, using REML, the mean 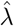 across 100 subsets was 0.85 (vs 0.95) under the network.

In simulations, 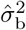 tended to be substantially larger when within-species variation was ignored, in which case 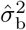 needs to compound between- and within-species variation. For leaflet length however, modeling versus ignoring within-species variation had little impact on 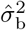. This again may be due to the large number of specimens. Using REML, when only 3 specimens were subsampled per morph, the 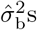 for 100 such subsets were on average about 9% larger (9.48% for the tree, 8.54% for the network) when within-species variation was ignored.

To fit BM_pheno_ and assess the impact of assuming equal phenotypic and phylogenetic relationships, we needed to reduce the data to specimens with both leaflet size and geographic data (997 out of 1757). Reducing the data had little effect on estimates and conclusions using BM_y_ (Table 3 for the full data, Table 4 for the reduced data). Using BM_pheno_, however, had a quite drastic effect compared to using BM_y_. The coefficient for elevation estimated using BM_pheno_ was still significantly negative, but much smaller in magnitude (Table 4). More importantly, BM_pheno_ estimates latitude to correlate positively with leaflet length, with strong evidence on the network and weak evidence on the tree. This is in contradiction with the negative correlation found using BM_y_.

**Table 4:**
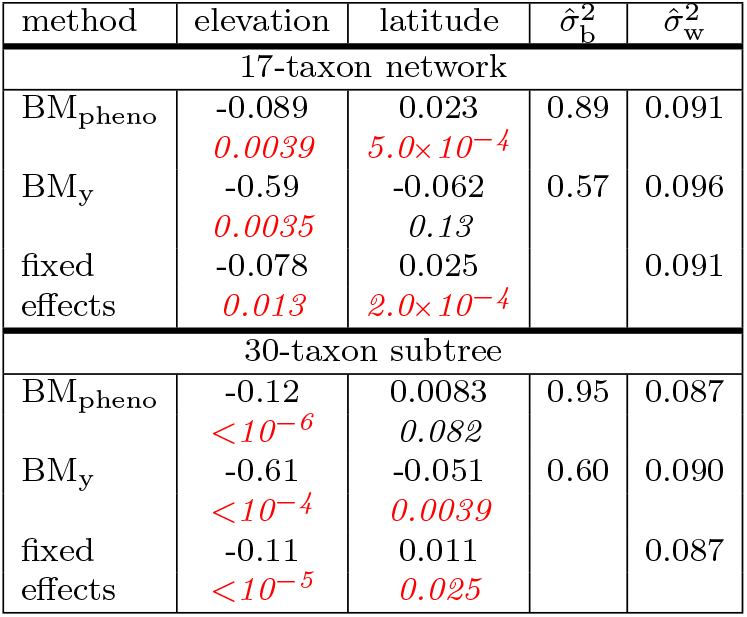
Analysis of leaflet length (log-transformed) with REML on the subset of specimens with both morphological and geographical data. BM_pheno_ assumes equal phenotypic and phylogenetic relationships. BM_y_ accounts for phenotypic variation in the response but not in the predictors. The model with fixed effects uses a standard linear regression on the individual-level data, with morph means (or intercept) estimated as fixed-effect parameters, and does not estimate 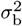. In the elevation and latitude columns, the first value is the estimated coefficient. The second italicized value is the associated p-value, in red when < 0.05.

This discordance suggests conflicting phenotypic and phylogenetic relationships. To estimate the phenotypic relationships alone, we fit a linear regression with morph means modeled as fixed effects (i.e. separate intercepts for each morph) and estimated as parameters, instead of estimating an evolutionary variance rate. We found a positive phenotypic coefficient for latitude, of magnitude greater than that estimated by BM_pheno_, on the network and on the tree (Table 4). This behavior matches our simulations under a phenotypic coefficient opposite to the evolutionary coefficient, in which the BM_pheno_ estimate was heavily biased towards the phenotypic coefficient, increasingly so with more specimens. The BM_y_ estimate was only slightly biased, and less so with more specimens. Therefore, we consider the BM_pheno_ estimates to be misleading for studying the evolution of leaflet size in *Polemonium*, because the phenotypic and evolutionary coefficients for latitude appear to be opposite, strongly violating the BM_pheno_ assumption.

#### 3.2.4 Leaflet size reconstruction

The predicted true mean of log leaflet length was obtained for *eddyense* (3 specimens) and *foliosissimum* (274 specimens). For *eddyense*, the observed mean log leaflet length was –1.22. The true morph mean was predicted at –1. 16 and – 1. 18 with and without elevation and latitude as predictors, representing 5.9% and 4.3% increases from the observed mean on the original scale. The prediction intervals (both of width 0.78) encompassed the observed mean. For *foliosissimum*, the observed mean was 0.622. The predicted means were both very close (0.621, a 0.1% decrease on the original scale) and the prediction intervals were narrow (width < 0.085). This illustrates that predictions for species with smaller sample sizes are more influenced by other species’ data, and less certain. Regardless of sample size, model predictors had little influence on the predictions.

At internal nodes, the width of prediction intervals increased with age: from 1.07 for the hybrid node directly ancestral to *elusum*, 1.72 for its minor parent, to 2.05 for the root. This pattern of ancestral state uncertainty increasing with distance from the tips was already known on trees [Ané, 2008].

## 4 Discussion

We presented a method to account for within-species trait variation on phylogenetic networks, a task with a long history on trees, and whose importance has been stressed by many authors. We reiterate the importance of this for avoiding overestimating the evolutionary variance. Intuitively, ignoring within-species variation is compensated for by an inflated evolutionary variance rate.

### 4.1 Methodology

Our approach is the first to allow for both within-species trait variation and reticulation, and to estimate within-species variance simultaneously with other model parameters instead of considering within-species variances as known without error. Our method assumes equal variances within species, and is robust to a violation of this assumption based on simulations. This assumption is also made by the PMM when used to account for within-species variation. It is suggested by Ives et al. [2007] as an option when the sample size per species is small, via the estimation of a pooled variance. When traits are hard to measure experimentally, typical sample sizes per species are very low. Moen et al. [2022] highlight this challenge for studies of adaptation, and the advantage of assuming equal variances in this context. Future method development could consider relaxing the assumption of homogeneous within-species variance for species with many sampled individuals.

Our implementation currently assumes that each hybrid node has exactly two parent lineages in the network, but the method allows for polytomies where a node has three or more children. Our implementation is currently limited to the BM, although our theory applies to more complex evolutionary models for the evolutionary covariance ***V***, at the cost of optimizing extra parameters. For example, Pagel’s λ model would include an independent component at the species level, beyond the variation between individuals or measurement error. It would also be interesting to allow for separate evolutionary rates along different parts of the phylogeny. In this case, ***V*** depends on the different rates and their mapping along the phylogeny [O’Meara et al., 2006]. As more complex models are developed, conclusions about evolutionary rates and phylogenetic signal could rely on likelihood ratio tests, although these tests are approximate. Tests based on bootstrapping procedures could be a possible future development to perform more accurate model comparisons.

Our simulations highlight the advantages of using REML instead of ML, especially with models that have multiple variance parameters, or to answer 5 questions about character rate evolution. For *Polemonium* leaflet length for example, switching from ML to REML can sway the estimate of phylogenetic signal from 0 to 1. The advantages of REML are well known [Ives et al., 2007] yet many software tools use ML only (e.g. `geiger`, Pennell et al. [2014] or `phylolm`, Ho and Ané [2014]). Unfortunately, REML is typically not an option for non-Gaussian generalized linear models, such as for phylogenetic logistic regression.

### 4.2 Importance of gene flow for traits

As of now, methods to estimate species networks scale poorly with the number of taxa. To detect gene flow and represent reticulations in a network, most studies focus their questions on a subsample of 20 taxa or so, a scale that methods such as SNaQ and Phylonet-MPL can handle [Hejase and Liu, 2016]. Downstream comparative analyses then face a dilemma: should they use more taxa on an approximate phylogeny without reticulation, or fewer taxa on a more accurate representation of the group’s phylogeny? Our case study on *Polemonium* suggests that using more taxa is advantageous and more powerful, as the increase in data quantity (and signal) can be substantial, out-weighing the approximation to the phylogenetic co-variance using a tree phylogeny. A caveat is that model mis-specification caused by ignoring reticulation on a tree may decrease power or increase type-1 error. We hope that this dilemma will disappear as network inference methods improve.

At each reticulation, one may ask which parent contributed to a given trait. Was the trait value inherited solely from one of the parents? Our BM model assumes that both parents contributed, such that the trait value at the reticulate node is a weighted average from the two parent values. The weights are the inheritance proportions (*γ*): the proportion of genes inherited from each parent. This is a legitimate prior for quantitative traits that are controlled by many genes of small and additive effects. But at each reticulation, one may ask if this null model is adequate for the trait under study.

To this end, Bastide et al. [2018] proposed a test for transgressive evolution after reticulation. This test can readily be used with our method to account for within-species trait variation.

Another approach is to compare the network model with a tree model, in which we assume that a trait is inherited from a single parent only, although model choice would need to account for the large number of options (up to 2^*h*^) with an increasing number *h* of reticulations. More generally, one may seek to optimize the weights (7) of the two parents at each reticulation, to best match evidence from the trait data. Bastide [2017] took this approach. Optimizing all h inheritance parameters could be too many, so he used a single parameter to scale the weights of all major hybrid edges simultaneously. Even with this amortized inference strategy, simulations showed that a single continuous trait variable had low information about the inheritance weights at reticulations.

Our findings in *Polemonium* are consistent. For all measures of leaflet size, the network model with inheritance values from genetic data was preferred over a tree model, in which the trait was forced to be inherited from a single parent (corresponding to inheritance values set to 0 or 1). However, the preference for the network was only very slight: the morphological signal is consistent with the genetic signal, but tenuous.

Multiple continuous traits would need to be combined to estimate the morphological signal for gene flow. As for trees, combining morphological traits is complicated by trait correlations. This caveat is especially important if we want to assume that traits share a common signal of gene flow. The traits more likely to have been inherited together through gene flow are the traits that share a genetic basis or form an integrated morphological component, and can be highly correlated with each other.

The inheritance signal may be stronger from discrete traits than from continuous traits, if the discrete trait is evolving slowly enough for accurate ancestral reconstruction. For example, Karimi et al. [2020] found support that flower color was introgressed during the evolution of baobabs in Madagascar. It would be interesting to extend our method for within-species variation to the study of discrete characters.

### 4.3 Phenotypic correlation

Our simulations highlight an important bias affecting many widely-used methods, when there is within-species variation in the predictors. The regression coefficient describing the historical evolutionary relationships are pulled towards the phenotypic coefficients. This bias is traditionally named “attenuation” when variation in the predictor is solely due to measurement error, uncorrelated with the other sources of variation [Fuller, 1987]. This pull decreases as the within-species sampling effort increases, for methods ignoring within-species variation in predictors.

For these methods, within-species variation in predictors causes a complex bias in estimating evolutionary variance rates. If a phenotypic relationship is absent or opposite to the evolutionary relationship, then 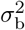 is overestimated. If the phenotypic relationship is equal or stronger than the evolutionary relationship, then 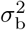 is underestimated. This interplay between phenotypic relationships (most often ignored for the study of long-term evolutionary patterns) and inference of evolutionary rates has not been identified before, to the best of our knowledge.

When predictors are available for the same set of individuals as the response trait, the BM_pheno_ model can be applied to account for within-species variation in predictors. However, BM_pheno_ assumes shared evolutionary and phenotypic relationships, such that the pull towards the phenotypic coefficients strengthens with more sampling effort, and the bias becomes extreme. We observed this for *Polemonium* leaflet size, where discordant evolutionary and phenotypic relationships led to opposite conclusions about the direction of correlation between leaflet length and latitude. For this reason, we recommend using this method with caution, and in combination with an assessment of the method’s assumption regarding phenotypic relationships. To estimate phenotypic correlations, standard linear models can be used with species as a fixed factor. Future work could tackle the question of rigorously testing whether phenotypic and evolutionary relationships are equal, extending the methods by Revell and Harmon [2008] and Goolsby et al. [2017] to reticulate phylogenetic networks and to a linear regression context (rather than correlation).

New methods are needed to handle the case when predictors are available on a different set of individuals than the response trait, if one wishes to use all individual values to best account for within-species trait variation, and to eliminate the pull of evolutionary coefficients towards phenotypic coefficients.

## Acknowledgements

We thank Cathy Cao for technical help setting up simulations. We also thank Joe Felsenstein and two anonymous reviewers for their thorough feedback, which helped improve the structure and clarity.

## Software and Data Availability

Our method is implemented in the `Julia` package `PhyloNetworks` available at https://github.com/crsl4/PhyloNetworks.jl starting with v0.14.0. Data and code for all simulations and analyses are available from the Dryad Digital Repository: https://doi.org/10.5061/dryad.9ghx3ffkc.

## Appendix

## A Parameter estimation

We prove here that (4) can be simplified to (5). Intuitively, (5) comes from reducing the model to species averages. The formula for 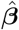 in (4) involves the inverse of ***W***_*η*_, so we first show that this large *N* × *N* matrix can in fact be inverted using smaller *n* × *n* matrices. Similar developments have reduced the computation complexity of some classes of mixed models [Demidenko, 2004, section 2.2.3], sometimes referred to as Henderson’s formula (1959). Our framework differs due to the phylogenetic correlation between species (“clusters” in the classical context). To express 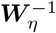, we apply the Woodbury matrix identity^5^:

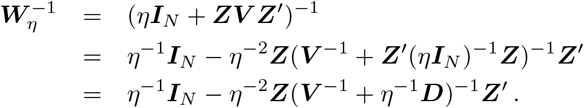

To invert ***V***^-1^ + *η*^-1^***D***, we apply the Woodbury matrix identity to *η****I***_*n*_ + ***VD***:

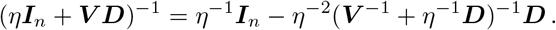

Using ***V***_*η*_ from (6), ***V***_*η*_***D*** = *η****I***_*n*_ + ***V D*** and we get:

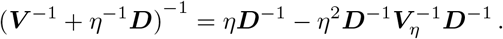

Combining the above equations, we get

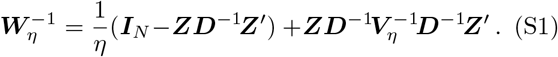

We are now ready to simplify (4), recalled here:

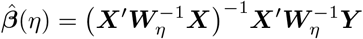

where ***X*** = ***Zx*** and 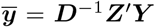. Using (S1) and ***Z′Z*** = ***D***, we have:

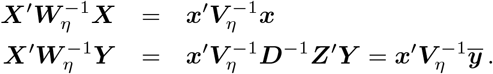

Combining the above equations gives (5).

We now turn to simplifying the profile likelihood criterion to be maximized, for the estimation of variance parameters (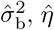 and any parameters for ***V***), to prove (7), (8) and (9). As usual, we instead write and seek to minimize twice the negative log (restricted) likelihood, denoted as 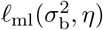 for ML and 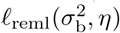 for REML:

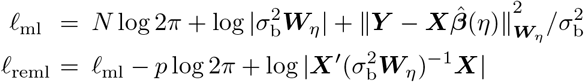

where we recall that 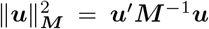. We now show how each term involving ***W***_*η*_ can be simplified using smaller matrices.

First, we use Sylvester’s determinant identity^6^ to express |***W***_*η*_| in terms |***V***_*η*_|.

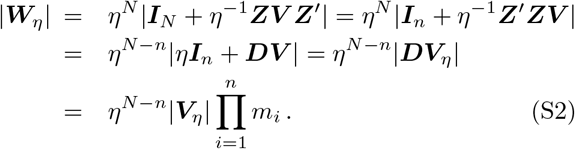

Next we use (S1) and (5) to simplify 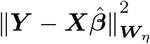.

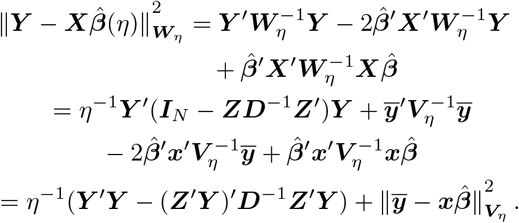

Recalling that 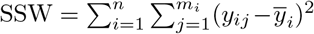 captures the sum of squared residuals within species, we get:

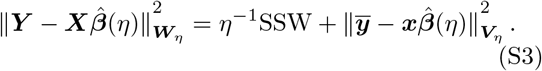

Next, we optimize 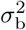 analytically as a function of *η* to profile ℓ_ml_ and ℓ_reml_ as functions of *η* only. If we fix *η* (and any other potential parameters for ***V***) and substitute (S2) and (S3) into ℓ_*ml*_ and ℓ_reml_, then we obtain the optimal value for 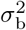 given in (9), which depends on the criterion via the degree of freedom *d* = *N* for ML and *d* = *N* – *p* for REML. Plugging 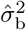 from (9) into ℓ_ml_ and ℓ_reml_ above, we obtain the profiled ML and REML criteria given in (7) and (8).

## B Parameter inference

To test hypotheses about a coefficient *β*_k_, we use its estimated standard error SE_k_ with

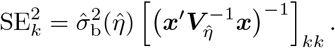

If the true *η* were known and used in the definition of SE_*k*_, then 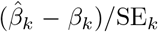 would follow a T-distribution with *N* – *p* degrees of freedom. But *η* is unknown. We approximate the distribution of 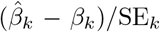 by a T-distribution with *n* – *p* degrees of freedom, being conservative by taking into account the number of species instead of the total number of observations. This approximation is exact in some classical contexts with balanced experiments, such as for the estimation of a population mean from *n* samples, each with *m* subsamples. More generally, a similar approximation is used for mixed models and has been shown to be superior to likelihood ratio tests for fixed effects [see e.g. Section 2.2.4 in Pinheiro and Bates, 2006]. Confidence intervals for regression co-efficients also use this approximation, assuming *n* – *p* degrees of freedom associated with 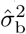.

## C Predicting species means

This task is traditionally called “ancestral state reconstruction”, but we favor the term “prediction”, as this task can be applied to present-day species. We use here the notations from section 2.3. In particular, ***y***_0_ denotes the true mean of species for which prediction is sought.

### C.1 Prediction variance

Section 2.3 gives the conditional mean ***μ***_0_(***β***) and variance **∑** of ***y***_0_ given 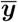, in the case when ***β*** is known. When ***β*** is estimated from 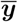, then the best prediction is 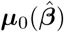 but has variance larger than **∑**. Namely, the prediction variance 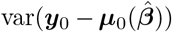 is then given by:

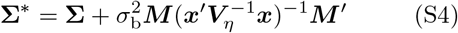

where ***M*** = ***x***_0_ – ***V***_*c*_***V***_*η*_^-1^***x*** [Christensen, 2001].

One might ask if conditioning on the individual-level data ***Y*** provides more information about ***y***_0_ than can be gained from the taxon-level means 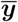. We show that both reduce to the same estimator, so that 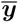 is sufficient for predictive purposes:

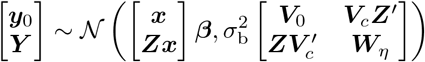

By applying (S1) and ***Z′Z*** = ***D*** we have:

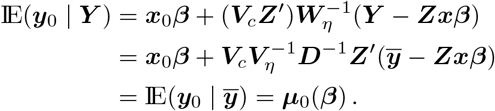

### C.2 Prediction interval

The prediction error 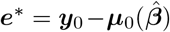 has distribution 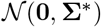 with **∑**\* given in (S4), so the error 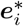 for the *i*th species to be predicted satisfies

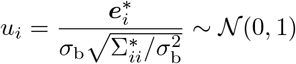

Note that the formula for 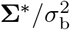 involves *η* but not 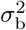. If *η* is known, then 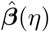 and 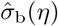 are independent by Cochran’s theorem. However, 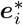 is not guaranteed to be independent of 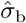. Nevertheless, we may use 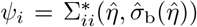 to estimate the variance of 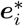 We then approximate the distribution of 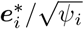 by a T-distribution with *n* – *p* degrees of freedom, as done above for testing fixed coefficients about between-species relationships.

Consequently, to build a 100(1 – *α*)% prediction interval for the *i*th species mean, we first find the (1 – *α*/2) quantile *q* of the T-distribution with *n* – *p* degrees of freedom and then use

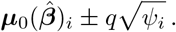

Recall here that formula (S4) for **∑**\* (hence 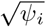) was obtained assuming that we know *η* and any other parameters for ***V***, and then simply plugging in their estimates in (S4). Doing so does not account for the extra uncertainty due to estimating *η*, which is hard to quantify [Christensen, 2001]. Hence, the prediction interval above should be considered as liberal.

### C.3 Example: Influence of other species information on reconstructed mean

We consider here the task of predicting the true mean for a species for which we do have data. In a simple example, we show that the prediction 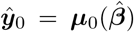 can be different from the observed species mean 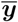, especially if a species has few sampled individuals. This example provides an intuition for what affects the prediction.

Suppose we have three taxa with sample sizes *m*_1_, *m*_2_, *m*_3_, and that the unscaled covariance matrix constructed from their phylogeny is:

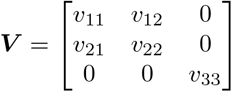

with taxon 1 and 2 sister to each other. Applying equations from section 2.3 and focusing on predicting the means for species 1 and 3, we get that

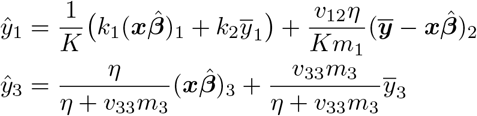

where 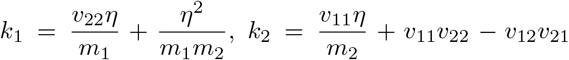 and *K* = *k*_1_ + *k*_2_. Therefore, 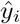 does not necessarily equal the sample mean 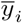. Instead, 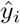 is pulled towards 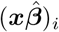, which depends on data across all species, and represents the ancestral state at the root if ***x*** is reduced to the intercept only. The pull is strong if *m_i_* is small or if within-species variation (*η*) is large. If *m_i_* is large, then the pull disappears and 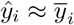.

Beyond the weighted average of 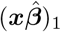 and 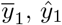 has an additional term proportional to the residual of its sister species 2. This term shows how information from closely related species is borrowed to influence prediction, in this simple example. As expected, this term vanishes when m1 increases.

## D Phenotypic correlation model

We study here the simulation model described in section 2.5, in which the within-species (or phenotypic) relationship between the response and the predictor traits differs from the between-species (or phylogenetic) relationship. We derive the distribution of the full response data ***Y*** and of the species means 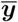 conditional on the observed predictor’s species means 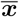. Since all variables are Gaussian, we simply need to derive the conditional means and variances. To do so, we repeatedly use the standard conditional distribution formulae for Gaussian processes.

Using (12) to simulate the predictor, (13) to simulate the response, the expression 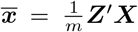 for species means, and the fact that 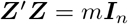, we get

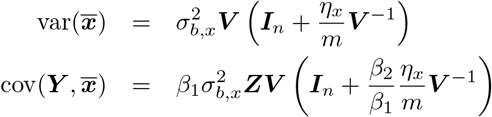

where 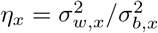. Therefore

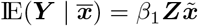

where we further define *u* = *η_x_/m* and

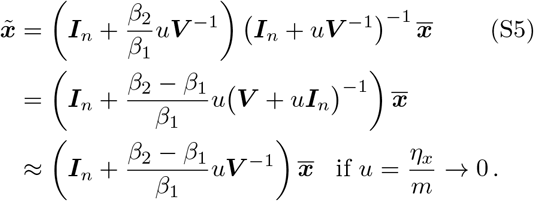

If *α*_1_ = *α*_2_ or 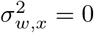 or *m* → ∞, then this simplifies to 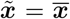, so that 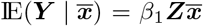 as assumed by our estimation model.

Next, 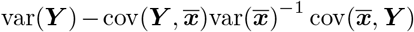 gives us

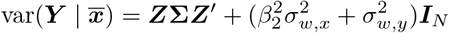

where

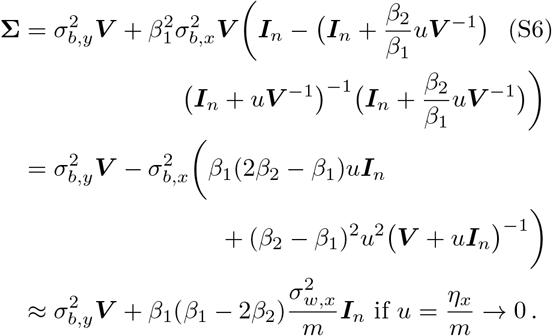

If 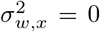 or *m* → ∞, then **∑** simplifies to 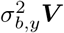 and the residual variance 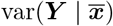 is as assumed in our estimation model.

For methods that ignore within-species variation, the conditional distribution of 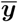 is relevant. From 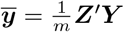 and our results above, we get

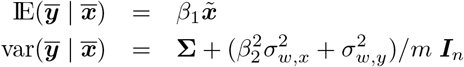

where 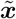 is as in (S5) and **∑** is as in (S6).

## E *Polemonium* leaflet analyses

**Figure S1:**
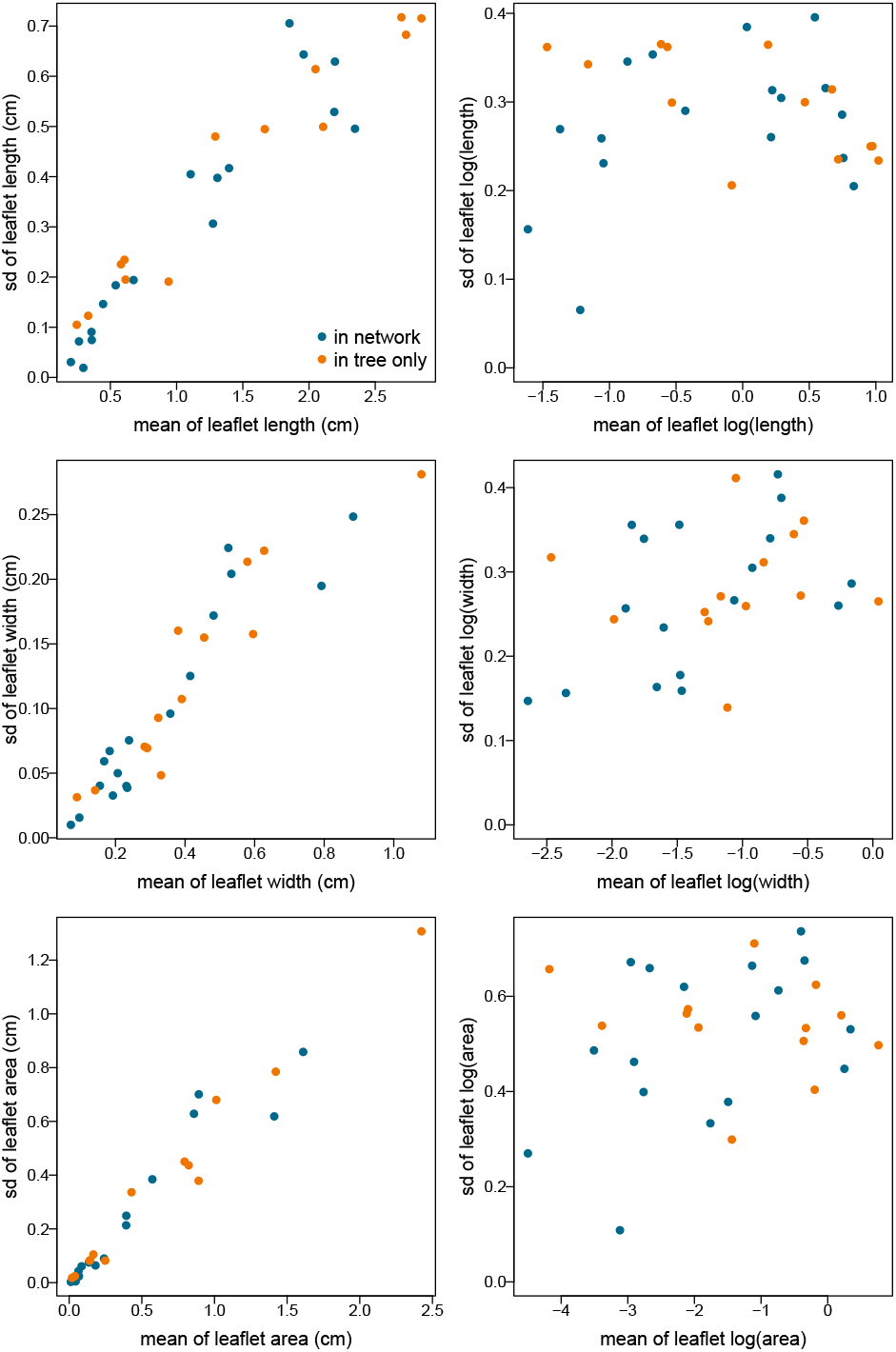
Log-transforming the response stabilizes within-morph variation and decorrelates it from mean response. The sample standard deviation (SD) in leaflet size is positively correlated with mean leaflet size across morphs (left), but not after transformation with the natural log (right). The spread of sample SDs also becomes more compact, reflecting a decrease in the relative variation of sample SDs across morphs.

**Figure S2:**
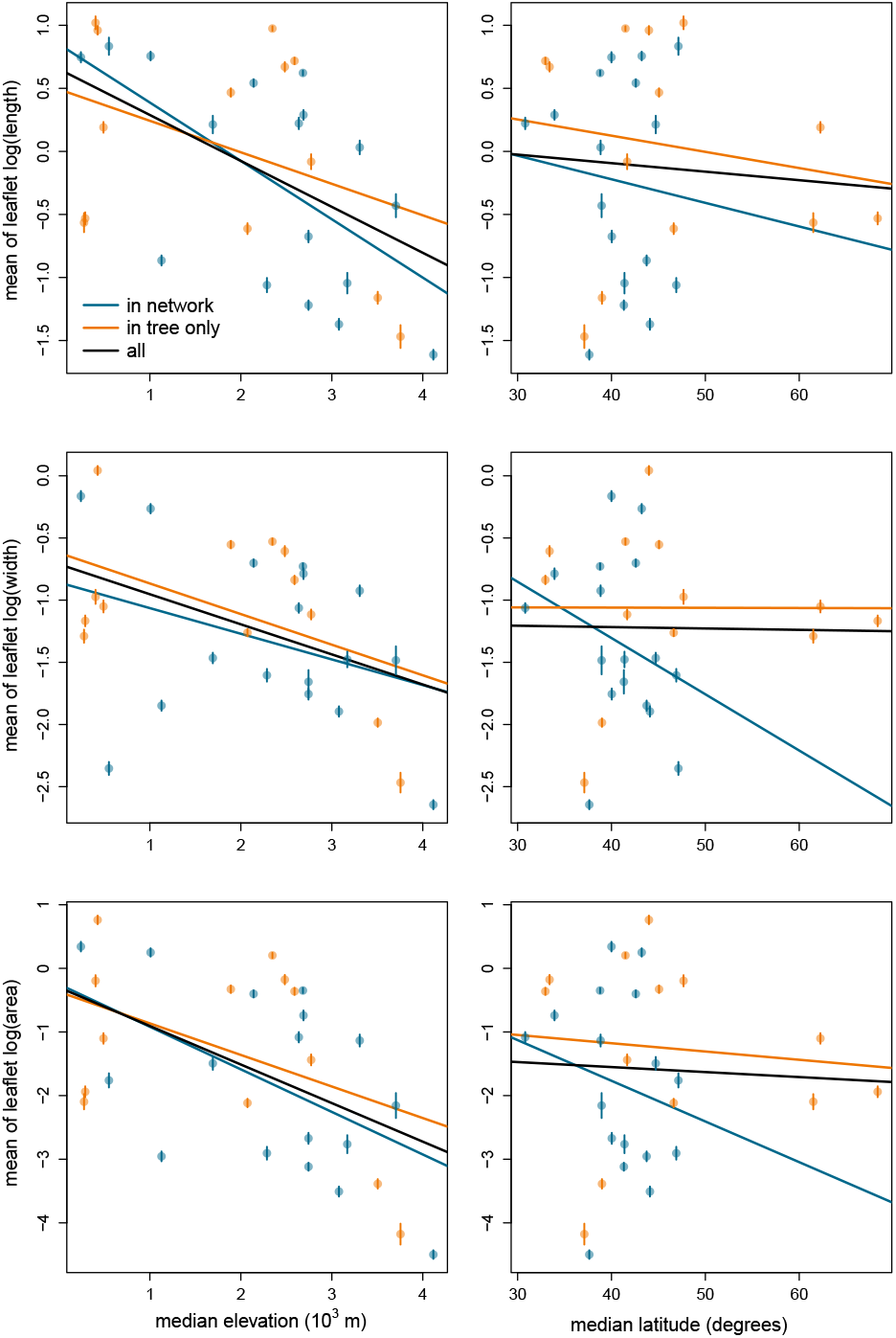
Leaflet size (log-transformed) versus elevation (left) and latitude (right). Each point represents a different morph. Colors indicate sampling across phylogenies: morphs in orange are in both the network and the tree. Morphs in blue are only in the tree. Vertical lines show ±1 standard error. The lines are based on ordinary (non-phylogenetic) simple linear regression using a single predictor, and either the orange or blue points only (orange and blue lines) or all points (black line).

**Table S1:**
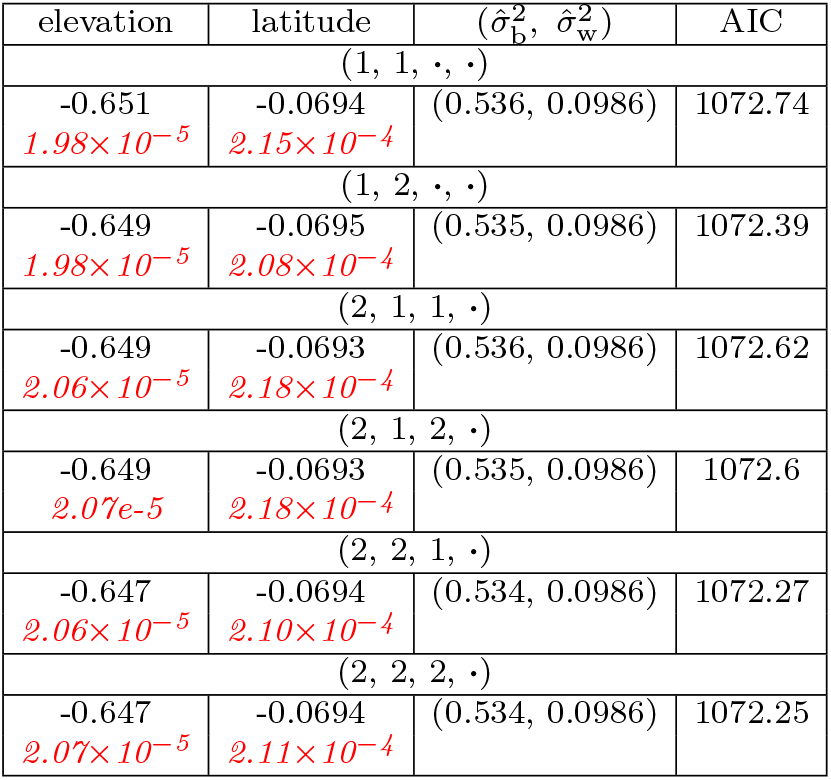
Results from fitting BM_y_ with REML on the 16 subtrees for log leaflet length. The subtrees are partitioned into 6 groups. Results are identical for all subtrees within the same group. Each group is represented by a 4-tuple, in which the first element is 1 (resp. 2) if *eximium* (resp. *eximium 2*) is selected; the second element is 1 (resp. 2) if *pulcherrimum p*. (resp. *pulcherrimum p. 2*) is selected, and the third element is 1 (resp. 2) if *chartaceum* (resp. *chartaceum 2*) is selected. The last element corresponds to the choice of the *californicum* accession. As it did not affect the results, both choices 1 and 2 are grouped and are represented by a dot. The last group (2, 2, 2, ·) was used in Tables 1, 3 and 4.

## Footnotes

1 https://intermountainbiota.org/portal/

2 https://www.pnwherbaria.org/

3 https://ucjeps.berkeley.edu/consortium/

4 *e*-0.908531 ≈ 0.40, *e*^-0.589785^ ≈ 0.55, *e*-0.319361 ≈ 0.73.

5 (***A*** + ***UCV***)^-1^ = ***A***^-1^(***I*** – ***U***(***C***^-1^ + ***V A***^-1^ ***U***)^-1^ ***V A***^-1^).

6 If ***A*** is *m*×*n* and ***B*** is *n*×*m*, then |***I***_*m*_ + ***AB***| = |***I***_*n*_ + ***BA***|.

## References

H. Akaike. A new look at the statistical model identification. IEEE Transactions on Automatic Control, 19(6):716–723, 1974. doi: 10.1109/TAC.1974.1100705.

C. Ané. Analysis of comparative data with hierarchical autocorrelation. The Annals of Applied Statistics, 2(3):1078–1102, 2008. doi: 10.1214/08-AOAS173.

P. Bastide. Shifted stochastic processes evolving on trees: application to models of adaptive evolution on phylogenies. Theses, Université Paris Saclay (COmUE), Oct. 2017. URL https://tel.archives-ouvertes.fr/tel-01629648.

P. Bastide, C. Solís-Lemus, R. Kriebel, K. William Sparks, and C. Ané. Phylogenetic comparative methods on phylogenetic networks with reticulations. Systematic Biology, 67(5): 800–820, 2018. doi: 10.1093/sysbio/syy033.

R. Christensen. Linear models for spatial data: Kriging, pages 269–311. Springer New York, New York, NY, 2001. ISBN 978-1-4757-3847-6. doi: 10.1007/978-1-4757-3847-6_6.

N. Cooper, G. H. Thomas, C. Venditti, A. Meade, and R. P. Freckleton. A cautionary note on the use of Ornstein Uhlenbeck models in macroevolutionary studies. Biological Journal of the Linnean Society, 118(1):64–77, 2016. doi: 10.1111/bij.12701.

E. Demidenko. Mixed Models: Theory and Applications. John Wiley & Sons, Inc., 2004. doi: 10.1002/0471728438.

J. Felsenstein. Phylogenies and quantitative characters. Annual Review of Ecology and Systematics, 19(1): 445–471, 1988. doi: 10.1146/annurev.es.19.110188.002305.

J. Felsenstein. Comparative methods with sampling error and within-species variation: Contrasts revisited and revised. The American Naturalist, 171(6): 713–725, 2008. doi: 10.1086/587525.

S. E. Fick and R. J. Hijmans. Worldclim 2: new 1-km spatial resolution climate surfaces for global land areas. International Journal of Climatology, 37(12): 4302–4315, 2017. doi: 10.1002/joc.5086.

W. A. Fuller. A single explanatory variable, pages 1–99. John Wiley & Sons, 1987. ISBN 978-0-4718-6187-4. doi: 10.1002/978-0-4703-1666-5.

L. Z. Garamszegi. Uncertainties due to within-species variation in comparative studies: measurement errors and statistical weights. In L. Z. Garamszegi, editor, Modern phylogenetic comparative methods and their application in evolutionary biology: concepts and practice, pages 157–199. Springer Berlin Heidelberg, Berlin, Heidelberg, 2014. ISBN 978-3-662-43550-2. doi: 10.1007/978-3-662-43550-2_7.

E. W. Goolsby, J. Bruggeman, and C. Ané. Rphylopars: fast multivariate phylogenetic comparative methods for missing data and within-species variation. Methods in Ecology and Evolution, 8(1):22–27, 2017. doi: 10.1111/2041-210X.12612.

T. F. Hansen and E. P. Martins. Translating between microevolutionary process and macroevolutionary patterns: The correlation structure of interspecific data. Evolution, 50(4):1404–1417, 1996. doi: 10.1111/j.1558-5646.1996.tb03914.x.

T. F. Hansen, J. Pienaar, and S. H. Orzack. A comparative method for studying adaptation to a randomly evolving environment. Evolution, 62(8): 1965–1977, 2008. doi: 10.1111/j.1558-5646.2008.00412.x.

L. Harmon. Phylogenetic comparative methods. Luke J. Harmon, 2019. URL https://lukejharmon.github.io/pcm/.

L. J. Harmon and J. B. Losos. The effect of intraspecific sample size on type i and type ii error rates in comparative studies. Evolution, 59(12):2705–2710, 2005. doi: 10.1111/j.0014-3820.2005.tb00981.x.

D. A. Harville. Bayesian inference for variance components using only error contrasts. Biometrika, 61 (2):383–385, 1974. doi: 10.1093/biomet/61.2.383.

H. A. Hejase and K. J. Liu. A scalability study of phylogenetic network inference methods using empirical datasets and simulations involving a single reticulation. BMC Bioinformatics, 17(1):422, 2016. doi: 10.1186/s12859-016-1277-1.

C. R. Henderson, O. Kempthorne, S. R. Searle, and C. M. von Krosigk. The estimation of environmental and genetic trends from records subject to culling. Biometrics, 15(2):192, June 1959. doi: 10.2307/2527669.

R. J. Hijmans. raster: geographic data analysis and modeling, 2020. URL https://cran.r-project.org/package=raster. R package version 3.4-5.

L. S. T. Ho and C. Ané. A linear-time algorithm for gaussian and non-gaussian trait evolution models. Systematic Biology, 63(3):397–408, 2014. doi: 10.1093/sysbio/syu005.

E. A. Housworth, E. P. Martins, and M. Lynch. The phylogenetic mixed model. The American Naturalist, 163(1):84–96, 2004. doi: 10.1086/380570.

J. Huang, Y. Thawornwattana, T. Flouri, J. Mallet, and Z. Yang. Inference of gene flow between species under misspecified models. Molecular Biology and Evolution, 39(12), 2022. doi: 10.1093/molbev/msac237.

A. R. Ives, P. E. Midford, and J. Garland, Theodore. Within-species variation and measurement error in phylogenetic comparative methods. Systematic Biology, 56(2):252–270, 2007. doi: 10.1080/10635150701313830.

N. Karimi, C. E. Grover, J. P. Gallagher, J. F. Wen-del, C. Ané, and D. A. Baum. Reticulate evolution helps explain apparent homoplasy in floral biology and pollination in baobabs (*Adansonia*; bom-bacoideae; malvaceae). Systematic Biology, 69(3): 462–478, 2020. doi: 10.1093/sysbio/syz073.

C. Körner, M. Neumayer, S. P. Menendez-Riedl, and A. Smeets-Scheel. Functional morphology of mountain plants. Flora, 182(5):353–383, 1989. ISSN 0367-2530. doi: 10.1016/S0367-2530(17)30426-7.

A. Kuznetsova, P. B. Brockhoff, and R. H. B. Christensen. lmerTest Package: Tests in Linear Mixed Effects Models. Journal of Statistical Software, 82: 1–26, 2017. doi: 10.18637/jss.v082.i13.

L. R. LaMotte. A direct derivation of the reml likelihood function. Statistical Papers, 48(2): 321–327, 2007. doi: 10.1007/s00362-006-0335-6.

G. E. Leventhal and S. Bonhoeffer. Potential Pitfalls in Estimating Viral Load Heritability. Trends in Microbiology, 24(9):687–698, 2016. doi: 10.1016/j.tim.2016.04.008.

M. Lynch. Methods for the Analysis of Comparative Data in Evolutionary Biology. Evolution, 45(5): 1065–1080, 1991. doi: 10.1111/j.1558-5646.1991.tb04375.x.

E. P. Martins and T. F. Hansen. Phylogenies and the comparative method: a general approach to incorporating phylogenetic information into the analysis of interspecific data. The American Naturalist, 149 (4):646–667, 1997.

D. S. Moen, E. Cabrera-Guzmán, I. W. Caviedes-Solis, E. González-Bernal, and A. R. Hanna. Phylogenetic analysis of adaptation in comparative physiology and biomechanics: overview and a case study of thermal physiology in treefrogs. Journal of Experimental Biology, 225(Suppl.1), 2022. doi: 10.1242/jeb.243292.

B. C. O’Meara, C. Ané, M. J. Sanderson, and P. C. Wainwright. Testing for different rates of continuous trait evolution using likelihood. Evolution, 60 (5):922–933, 2006. doi: 10.1554/05-130.1.

M. Pagel. Inferring the historical patterns of biological evolution. Nature, 401(6756): 877–884, 1999. doi: 10.1038/44766.

H. D. Patterson and R. Thompson. Recovery of inter-block information when block sizes are un-equal. Biometrika, 58(3): 545–554, 1971. doi: 10.1093/biomet/58.3.545.

M. Pennell, J. Eastman, G. Slater, J. Brown, J. Uyeda, R. Fitzjohn, M. Alfaro, and L. Harmon. geiger v2.0: an expanded suite of methods for fitting macroevolutionary models to phylogenetic trees. Bioinformatics, 30:2216–2218, 2014.

J. Pinheiro and D. Bates. Mixed-Effects Models in S and S-PLUS. Springer Science & Business Media, 2006. ISBN 9780387227474.

L. J. Revell. phytools: An R package for phylogenetic comparative biology (and other things). Methods in Ecology and Evolution, 3:217–223, 2012.

L. J. Revell and L. J. Harmon. Testing quantitative genetic hypotheses about the evolutionary rate matrix for continuous characters. Evolutionary Ecology Research, 10:311–331, 2008.

J. P. Rose. Taxonomy and relationships within *Pole-monium foliosissimum* (Polemoniaceae): untangling a clade of colorful and gynodioecious herbs. Systematic Botany, 46(3):519–537, 2021. doi: 10.1600/036364421X16312067913372.

J. P. Rose, C. A. P. Toledo, E. M. Lemmon, A. R. Lemmon, and K. J. Sytsma. Out of sight, out of mind: widespread nuclear and plastid-nuclear discordance in the flowering plant genus *Polemonium* (Polemoniaceae) suggests widespread historical gene flow despite limited nuclear signal. Systematic Biology, 70(1):162–180, 2021. doi: 10.1093/sysbio/syaa049.

J. Schindelin, I. Arganda-Carreras, E. Frise, V. Kaynig, M. Longair, T. Pietzsch, S. Preibisch, C. Rueden, S. Saalfeld, B. Schmid, J.-Y. Tinevez, D. J. White, V. Hartenstein, K. Eliceiri, P. Tomancak, and A. Cardona. Fiji: an open-source platform for biological-image analysis. Nature Methods, 9(7): 676–682, 2012. doi: 10.1038/nmeth.2019.

D. Silvestro, A. Kostikova, G. Litsios, P. B. Pearman, and N. Salamin. Measurement errors should always be incorporated in phylogenetic comparative analysis. Methods in Ecology and Evolution, 6 (3):340–346, 2015. doi: 10.1111/2041-210X.12337.

C. Solís-Lemus, P. Bastide, and C. Ané. Phylonetworks: a package for phylogenetic networks. Molecular Biology and Evolution, 34(12): 3292–3298, 2017. doi: 10.1093/molbev/msx235.

I. J. Wright, N. Dong, V. Maire, I. C. Prentice, M. Westoby, S. Díaz, R. V. Gallagher, B. F. Jacobs, R. Kooyman, E. A. Law, M. R. Leishman, Ü. Niinemets, P. B. Reich, L. Sack, R. Villar, H. Wang, and P. Wilf. Global climatic drivers of leaf size. Science, 357(6354): 917–921, 2017. doi: 10.1126/science.aal4760.

